# An inducible AraC that responds to blue light instead of arabinose

**DOI:** 10.1101/2020.07.14.202911

**Authors:** Edoardo Romano, Armin Baumschlager, Emir Bora Akmeriç, Navaneethan Palanisamy, Moustafa Houmani, Gregor Schmidt, Mehmet Ali Öztürk, Leonard Ernst, Mustafa Khammash, Barbara Di Ventura

## Abstract

In *Escherichia coli*, the operon responsible for the catabolism of L-arabinose is regulated by the dimeric DNA-binding protein AraC. In the absence of L-arabinose, AraC binds to the distal I_1_ and O_2_ half-sites, leading to repression of the downstream P_BAD_ promoter. In the presence of the sugar, the dimer changes conformation and binds to the adjacent I_1_ and I_2_ half-sites, resulting in the activation of P_BAD_. Here we engineer blue light-inducible AraC dimers in *Escherichia coli* (BLADE) by swapping the dimerization domain of AraC with blue light-inducible dimerization domains. Using BLADE to overexpress proteins important for cell shape and division site selection, we reversibly control cell morphology with light. We demonstrate the exquisite light responsiveness of BLADE by employing it to create bacteriographs with an unprecedented quality. We then employ it to perform a medium-throughput characterization of 39 *E. coli* genes with poorly defined or completely unknown function. Finally, we expand the initial library and create a whole family of BLADE transcription factors (TFs), which we characterize using a novel 96-well light induction setup. Since the P_BAD_ promoter is commonly used by microbiologists, we envisage that the BLADE TFs will bring the many advantages of optogenetic gene expression to the field of microbiology.

---

While the preferred carbon source for *E. coli* under most conditions is glucose, other sugars, such as lactose or arabinose, also support cell growth, albeit typically at a slower rate^1, 2^. Three operons are responsible for the uptake and catabolism of L-arabinose: the *BAD* operon, encoding three catabolic enzymes that convert L-arabinose to D-xyluluose-5-phosphate; the *FGH* operon, encoding the transporters that regulate L-arabinose uptake when its concentration in the extracellular environment is low, and the *araE* operon, encoding a low-affinity transporter that acts at high extracellular L-arabinose concentrations^3, 4^. In the absence of L-arabinose, P_BAD_ is repressed by AraC, the regulator of the system, bound to the distal I_1_ and O_2_ half-sites, which causes the formation of a DNA loop that sterically blocks the access of the RNA polymerase to the promoter (Fig. 1a). In the presence of L-arabinose, transcription from the P_BAD_, P_FGH_ and P_E_ promoters is activated by AraC, which additionally negatively feeds back on its own promoter P_C_, found upstream of, and in reverse orientation to, P_BAD_^3, 4^. Activation results from AraC binding to the adjacent I_1_ and I_2_ half-sites, which recruits the RNA polymerase (Fig. 1a). AraC is composed of an N-terminal dimerization domain (DD) and a C-terminal DNA binding domain (DBD) connected via a linker (Fig. 1b). Interestingly, AraC is always a homodimer, whether bound to arabinose or not^4^. Binding of arabinose triggers a conformational change in AraC, which results in the two DBDs being oriented in a way that favors their interaction with the I_1_ and I_2_ half-sites rather than the I_1_ and O_2_ half-sites (Fig. 1a)^3, 4^. The mechanism explaining this conformational change, which involves ligand-induced regulation of the position of the N-terminal arm of AraC, has been named the light switch, despite AraC not being a photoreceptor^3^. We reasoned that, if AraC could be made to respond to light, as previously done for other bacterial and eukaryotic transcriptional regulators^5, 6^, it would be possible for microbiologists to reversibly steer, with high spatio-temporal resolution, a great variety of biological processes relying on gene expression. They would simply employ well-known P_BAD_-based vectors, such as pBAD33, modified only to express the engineered light-sensitive AraC in place of the arabinose-sensitive natural one. Importantly, strains previously constructed to control with arabinose a genomic locus, in which the P_BAD_ promoter was inserted in place of an endogenous promoter^7–9^, would be fully compatible with this system. Here we show that, by swapping the AraC DD with the blue light-triggered dimerizing protein VVD^10^, and by selecting the appropriate linker between VVD and the DBD, we are able to render AraC blue light responsive. We characterize this novel AraC, which we name BLADE (for Blue Light-inducible AraC Dimers in *E. coli*), in terms of kinetics, reversibility, and light dependence. Taking advantage of the ability of BLADE to trigger gene expression only in illuminated cells, we perform bacterial photography and reproduce the Blade Runner movie poster at high resolution using a lawn of bacteria expressing the superfolder green fluorescent protein (sfGFP) under the control of BLADE. We then utilize BLADE to control *E. coli* cell morphology by overexpressing MinD^Δ10^, MreB and RodZ. Employing a previously constructed *E. coli* strain where endogenous *rodZ* is under the control of the P_BAD_ promoter^7^, we demonstrate that light, but not arabinose, allows for the reversible switching between round and rod cell morphologies. To showcase the advantage of light as external trigger in medium and high-throughput assays, we build a library of 117 constructs to characterize 39 *E. coli* genes with unknown or poorly defined function in terms of intracellular localization and effect on cell growth and morphology. We investigate the mechanism of BLADE action *in vivo*, and show that beyond contacting the I_2_ half-site in the lit state, the dark state involves the formation of aggregates, which likely contribute to the tightness of the system. We engineer an entire family of BLADE TFs creating a much larger library comparing two light-inducible dimerization domains, different linkers and positioning of the components. Interestingly, we find that the order of the light-dependent dimerizing and DBD domains does not need to resemble that of wild type AraC. We show that a synthetic promoter containing two I_1_ half-sites is still light-inducible and leads to higher light/dark fold change in gene expression compared to the wild type P_BAD_ promoter based on I_1_-I_2_ in a small range of BLADE concentrations. Finally, we develop a high-throughput characterization approach using a novel 96-well light induction setup, which can be easily built and employed, to find optimal expression levels of the BLADE TFs for best performance. We envision that BLADE will stimulate the incorporation of optogenetic experiments in microbiology due to its compatibility with previously constructed strains and plasmids, its added functionality that cannot be easily achieved with chemical inducers, and its reliable performance.

**Fig. 1.**
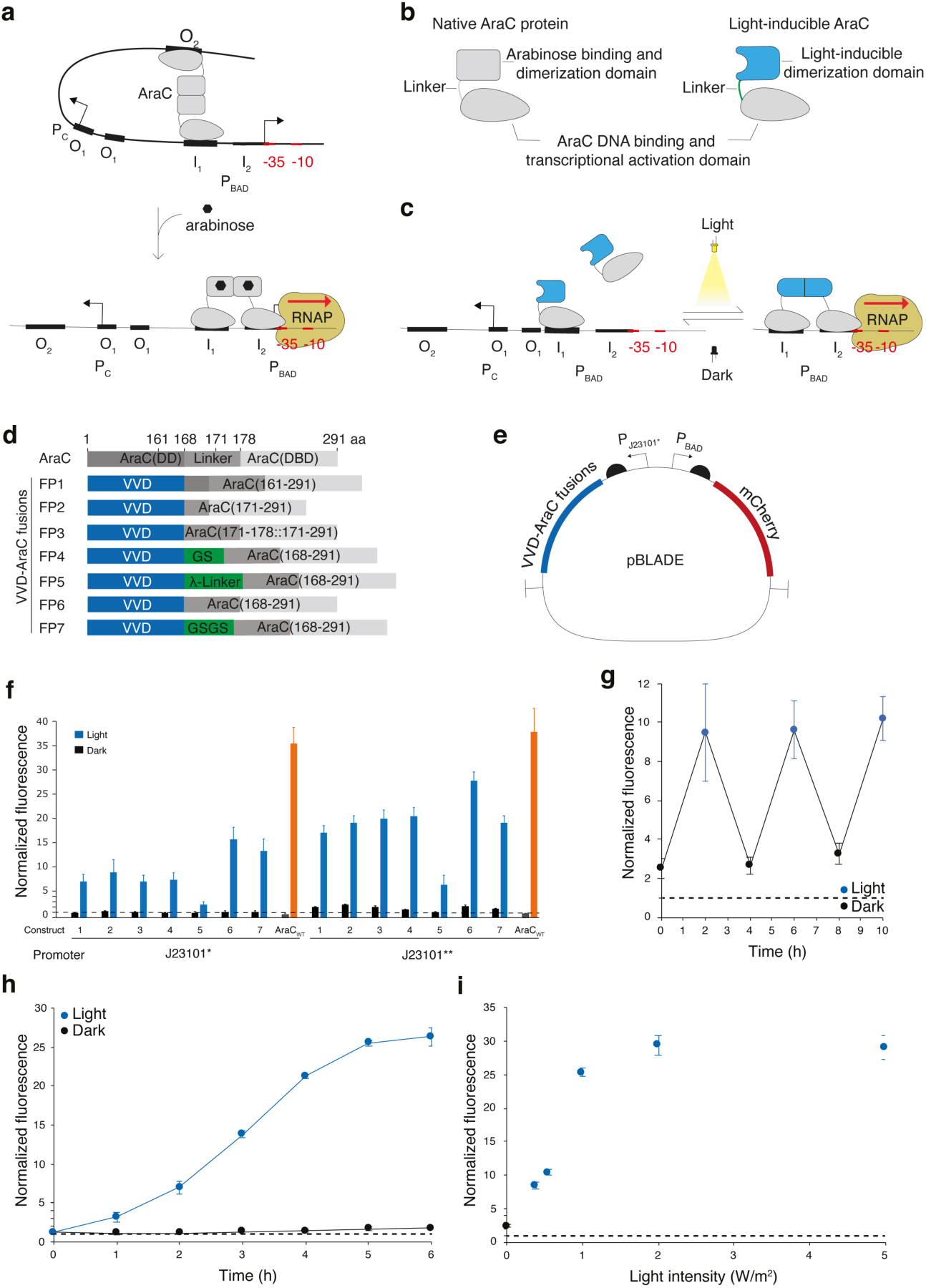
**Engineering and characterization of a novel light-inducible AraC a**, Mechanism of arabinose-induced P_BAD_ induction by AraC. The thickness of the I_1_ and I_2_ half-sites is proportional to the affinity with which AraC binds to them. P_C_, promoter driving the expression of *araC*. **b**, Domain composition of wild type (left) and light-inducible (right) AraC. **c**, Expected mechanism of P_BAD_ activation by light-inducible AraC. **d**, Domain composition of the chimeric VVD-AraC_DBD_ fusion constructs. In FP3, amino acids 171-178 of the natural linker are present twice. **e**, Plasmid for expression of a gene of interest (here mCherry) under control of BLADE. **f**, mCherry fluorescence intensity in *E. coli* MG1655 cells transformed with the library shown in (**d**) grown for 4h either in the dark or under 460 nm light (5 W/m^2^) illumination. Native AraC cloned under the same constitutive promoters was used as positive control. The bars for AraC represent the values obtained without (black) and with 0.1% (orange) arabinose for 4h. **g**, mCherry fluorescence intensity in the same culture of *E. coli* MG1655 cells transformed with the FP6 fusion driven by the J23101** promoter after repeated cycles of blue light exposure and darkness. **h**, Kinetics of mCherry expression in *E. coli* MG1655 cells transformed with the FP6 fusion driven by the J23101* promoter grown for 4h either in the dark or under 460 nm light (5 W/m^2^) illumination. **i**, mCherry fluorescence intensity measured in *E. coli* MG1655 cells transformed with the FP6 fusion driven by the J23101** promoter grown for 4h under 460 nm light of the indicated light intensity (cyan) or kept in the dark for 4h (black). **f-i**, All values were normalized to the mCherry fluorescence intensity measured in *E. coli* MG1655 cells transformed with the plasmid deprived of the transcription factor (dashed line). Values represent mean ± s.d. of at least *n=3* independent experiments.

## Results

### Creation of a small library of chimeric VVD-AraC fusion constructs

Inspired by a previous study in which chimeric AraC constructs have been cloned to probe the role of the DD and DBD^11^, we reasoned that, by exchanging the dimerization domain of AraC with a light-inducible dimerization domain (Fig. 1b), we would be able to control with light the switching of this engineered AraC from monomer to dimer (Fig. 1c). In its monomeric form, the engineered AraC would contact the high-affinity I_1_ half-site^12^, but not the low-affinity I_2_ half-site, needed to recruit the RNA polymerase. Its function as a light-inducible TF would depend on finding the appropriate linker supporting the correct orientation of the two DBDs after dimer formation, permissive of I_1_-I_2_ binding (Fig. 1c). As the light-triggered dimerization domain we selected VVD, which has often been successfully employed to control with light the dimerization of proteins of interest^5, 6, 13, 14^. VVD senses blue light *via* the flavin adenine di-nucleotide (FAD) chromophore^10^. Blue light triggers the formation of a cysteinyl-flavin adduct, which generates a new hydrogen bond network that releases the N-terminus (N-terminal cap) from the protein core and restructures it creating a new dimerization interface^15, 16^. We swapped AraC dimerization domain with VVD^N56K/C71V^, a double mutant shown to stabilize the dimer^5^, and cloned seven constructs having different linkers between AraC_DBD_ and VVD (Fig. 1d). We removed the *araC* gene from pBAD33, and introduced two constitutive promoters of different strength (J23101* and J23101**) to drive the expression of the chimeric VVD-AraC_DBD_ fusion constructs (Supplementary Fig. 1). For a reporter gene, we cloned *mCherry* downstream of the P_BAD_ promoter (Fig. 1e). As positive control, we constructed the same plasmid carrying full-length AraC in place of the VVD-AraC fusion (Supplementary Fig. 2c), while the plasmid without any TF was constructed to serve as negative control to monitor leaky expression from P_BAD_ (Supplementary Fig. 2b). Flow cytometry analysis of *E. coli* MG1655 cells transformed with the small library of VVD-AraC fusions, as well as the negative and the positive controls, kept in the dark or illuminated with 460 nm light (5 W/m^2^) for 4 hours showed that all 14 VVD-AraC constructs were light-inducible, despite being less optimal than full-length AraC (Fig. 1f and Supplementary Fig. 3). Different linkers corresponded to different amounts of gene expression. With the weaker constitutive promoter driving expression of the VVD-AraC_DBD_ fusion constructs (JS23101*), the levels of reporter expression in the dark approached those of the negative control, to which the values were normalized (Fig. 1f). The stronger constitutive promoter (JS23101**) led to significantly higher expression of the reporter gene after blue light illumination for all constructs, albeit at the cost of increased leakiness in the dark (Fig. 1f). Nonetheless, for some of the fusions, the light/dark fold change was higher with this promoter. We named a generic member of this family of Blue Light-inducible AraC Dimers in *E. coli* BLADE and the pBAD33-derived corresponding expression plasmid pBLADE (Fig. 1e). Before continuing with further characterization and utilization of BLADE, we verified that the blue light required for its activation was well tolerated by the cells (Supplementary Fig. 4).

### Characterization of BLADE

A clear advantage of light as an external trigger is that it can be easily turned off, enabling reversible control of gene expression, without the need of potentially damaging and time-consuming washing steps. We exposed *E. coli* MG1655 cells transformed with pBLADE-mCherry to alternating 2 hours-cycles of blue light and darkness, for a total of 3 illumination cycles and measured mCherry levels via flow cytometry at the end of every phase. The same expression levels were reached after every illumination phase, and the reporter was repressed to the same extent after every dark phase (Fig. 1g). Next, we measured the kinetics of mCherry expression from pBLADE and found that the half-maximum was reached after about 2.5 hours of induction with light, while the levels plateaued after 5 hours of light induction (Fig. 1h). To assess the requirement of BLADE in terms of blue light, we measured mCherry levels obtained at different light intensities, and found that 1 W/m^2^ was sufficient to obtain reporter gene expression levels close to the saturation value (Fig. 1i).

Next we sought to demonstrate that BLADE is useful to control the expression of functional *E. coli* proteins, and not only fluorescent reporters. We thus cloned *eyfp-minD* into pBLADE. MinD is a dynamically membrane-bound ATPase, which, together with MinC and MinE, constitutes the Min system, a machinery needed to place the divisome at mid-cell^17^ and to aid chromosome segregation^18^. In order to exert its inhibitory function against FtsZ, the protein that starts divisome assembly, MinC must be recruited to the cytoplasmic membrane by MinD^19, 20^. In *E. coli*, MinD – and, consequently, MinC – oscillate from pole to pole due to the action of MinE, with a period of 50-60 seconds at room temperature^21^. When averaged over time, MinC/D concentration is highest at the poles and minimal at mid-cell, causing the septum to form at mid-cell. Overexpression of MinD results in filamentation, because endogenous MinE is not sufficient to displace all MinD molecules from the membrane, allowing the MinCD complex to become stably and homogenously membrane-bound, inhibiting FtsZ everywhere^17^. Thus, MinD is a good candidate to check the tightness of the BLADE system. Time-lapse fluorescence microscopy showed that eYFP-MinD oscillations were present only in cells illuminated with blue light and not in those kept in the dark (Supplementary Fig. 5a and Supplementary Video 1). The distribution of the cell length for both non-induced and induced samples was comparable to that of the same strain transformed with the negative control (Supplementary Fig. 5b).

### Spatial control of gene expression

One of the benefits of optogenetic induction is the ability to modulate gene expression in a spatially dependent fashion. To showcase how BLADE could be used to control expression of a target gene only in selected cells, we cloned sfGFP^22^ into pBLADE. *E. coli* MG1655 cells transformed with pBLADE-sfGFP were then applied to an agar pad and subjected to confocal microscopy to expose a limited area (6.4 μm^2^) to blue light every 5 minutes. After 3 hours, sfGFP was expressed up to 6.7-fold more in the illuminated cells compared to the surrounding non-illuminated cells (Supplementary Fig. 6). Another interesting application of light-inducible TFs that relies on the possibility to shine light on a plate in desired patterns, is bacterial photography^23^. To assess the effectiveness of BLADE in this type of application, we covered a lawn of *E. coli* MG1655 cells transformed with pBLADE-sfGFP with a photomask depicting the Blade Runner movie poster (Fig. 2a). We illuminated the plate with blue light overnight and then took several microscopy pictures and stitched them together (Fig. 2b). The sensitive light response of BLADE yielded a good contrast, resulting in a high quality bacteriograph that allowed for the faithful reproduction of the details in the poster, such as facial expressions (Fig. 2c).

**Fig. 2.**
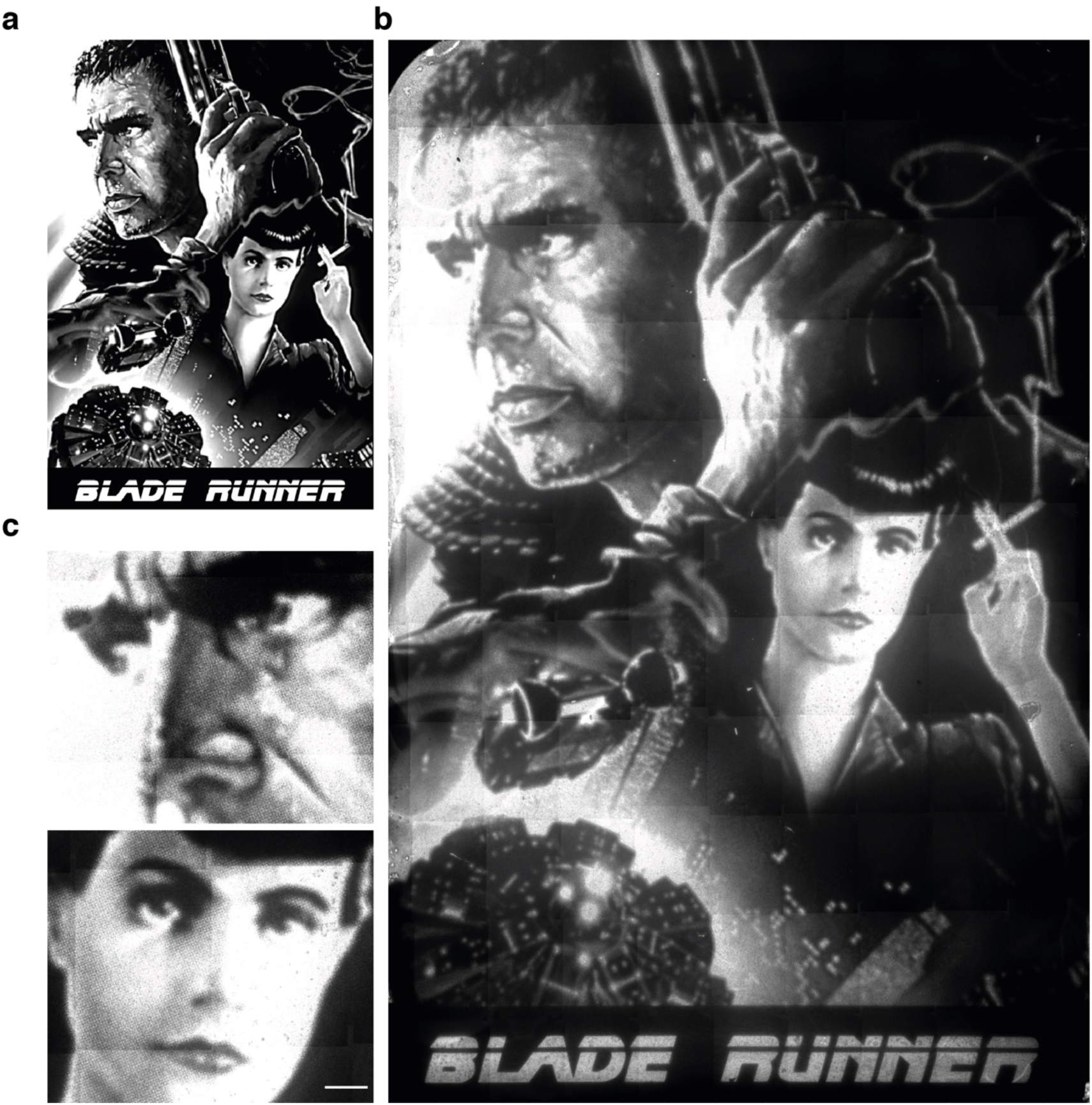
**BLADE allows for the production of high contrast bacteriographs. a**, Photomask used to produce the bacteriograph (printed with permission from Warner Bros. Entertainment, Inc.). **b**, Bacteriograph. A lawn of *E. coli* MG1655 cells transformed with pBLADE(FP6*)-sfGFP were grown overnight at 37°C while being exposed to blue light through the photosmask in (**a**). 110 individual images were taken with a fluorescent microscope and stitched together via image analysis software. Scale bar, 1 cm. **c**, Zoom in on two parts of the bacteriographs. Scale bar, 300 μm.

### Controlling *E. coli* cell morphology with BLADE

Cell morphology impacts growth and survival in diverse environments^24, 25^. Being able to generate desired cell morphologies with light could pave the way to performing experiments to better understand the contribution of cell morphology to bacterial fitness and adaptation to a particular environment. To demonstrate light control of cell morphology, we selected three *E. coli* proteins to overexpress: MinD^Δ10^, MreB and RodZ. MinD^Δ10^ is a truncated form of MinD lacking the last 10 amino acids constituting the membrane targeting sequence (MTS). Without the MTS, MinD^Δ10^ cannot associate with the membrane and remains cytoplasmic. It however maintains the ability to homodimerize^26^. We hypothesized that MinD^Δ10^ could heterodimerize with endogenous MinD. The heterodimer formed by MinD and MinD^Δ10^ would not be able to stably bind to the membrane, because a monovalent MTS is not sufficient for this^27^. With MinD sequestered into the cytoplasm, endogenous MinC would no longer be recruited to the membrane, and FtsZ should be free to start divisome assembly at the poles, leading to the formation of anucleated mini-cells and variably long cells (Fig. 3a), which would manifest as a multimodal cell length distribution in a population of cells. MreB is the bacterial actin homolog, necessary for the establishment and maintenance of rod shape and cell wall synthesis^28–30^. Its assembly is regulated by RodZ, a transmembrane protein that binds MreB, altering the conformational dynamics and intrinsic curvature of MreB polymers^31–33^. It has been previously established that overexpression of MreB or RodZ leads to cell elongation and thickening^30, 31, 34^. We cloned the above-mentioned genes into pBLADE and transformed each into MG1655 *E. coli* cells. We then exposed the cells to 4 hours of blue light illumination. Cells kept in the dark served as controls. BLADE-induced MinD^Δ10^ overexpression led to the formation of minicells; cells kept in the dark were indistinguishable from those transformed with an empty pBLADE, which served as negative control (Fig. 3b, c). The phenotype was not caused by the illumination (Fig. 3c). In contrast to MinD^Δ10^ overexpression, BLADE-induced MreB and RodZ overexpression led to cell elongation and thickening, while cells kept in the dark were indistinguishable from the negative control (Fig. 3d-f). We additionally controlled endogenous RodZ with BLADE using a previously constructed strain (KC717), where the endogenous promoter driving *rodZ* expression has been exchanged with P_BAD_^7^. In the absence of arabinose, the endogenous chromosomal copy of AraC inhibits transcription from P_BAD_, thus RodZ is not expressed and cells are spherical^7, 31, 32, 35^. In the presence of arabinose, endogenous AraC initiates transcription from P_BAD_ and, consequently, RodZ is expressed, leading to the reappearance of rod-shaped cells^7^. We transformed KC717 cells either with pBLADE-RodZ (population A) or with an empty pBAD33 deprived of *araC* and P_BAD_ (pBAD#; population B) and kept both populations either uninduced (in the dark for population A, and without arabinose for population B) or induced them for 4 hours (with blue light for population A and with arabinose for population B) (Fig. 3g). At this time point, population A recovered the rod-shape to a greater extent than population B (Fig. 3h). To showcase the power of optogenetics to quickly switch induction off, we subjected the cells to a recovery phase, by putting them into the dark (population A) and washing arabinose off (population B). While it was possible to obtain spherical cells again after 2 hours of dark incubation, the cells that had been induced with arabinose did not recover the initial phenotype and rather became even more rod-shaped (Fig. 3h).

**Fig. 3.**
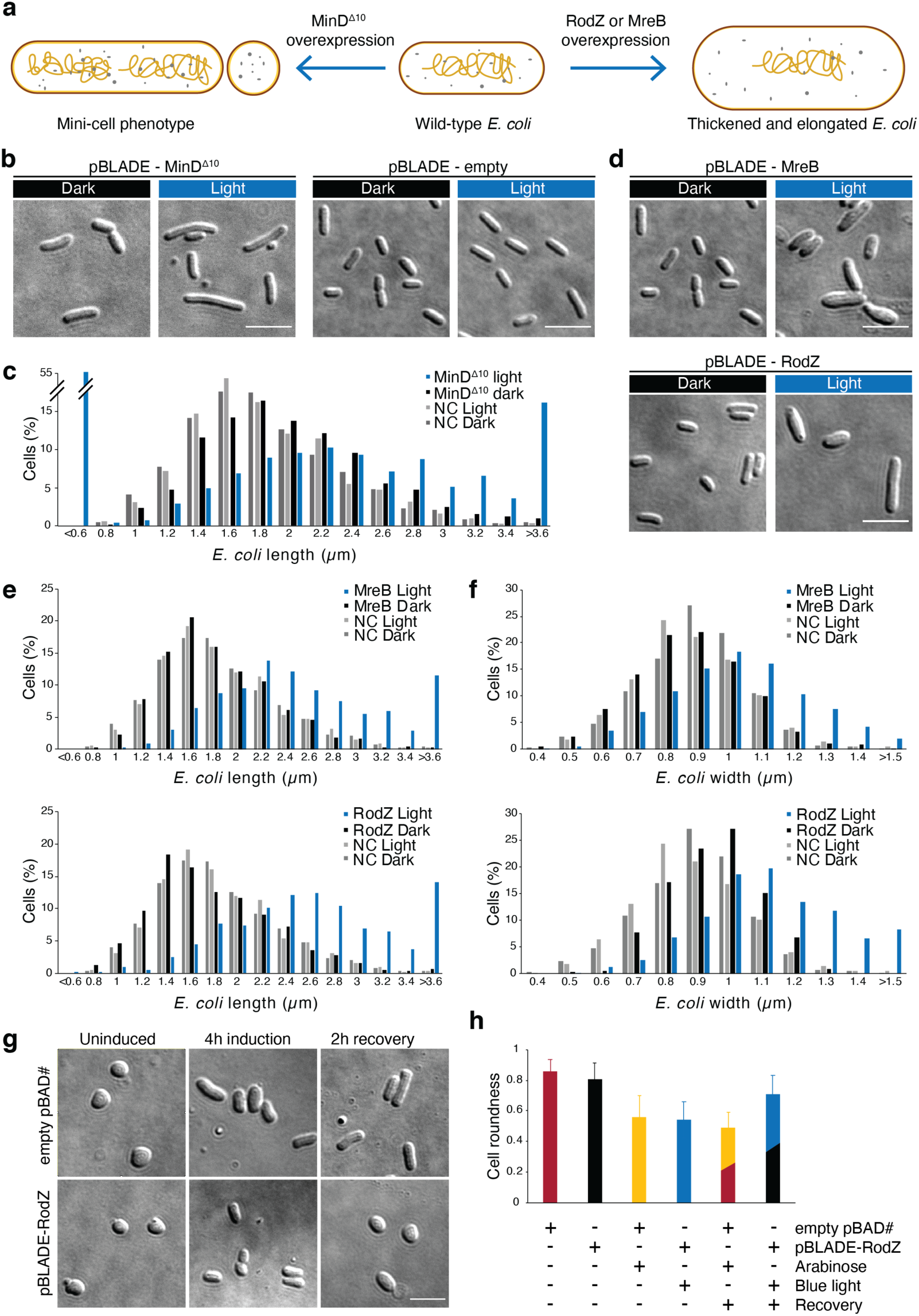
**BLADE enables reversible control of cell morphology. a**, Phenotypes expected when overexpressing the indicated proteins. **b**,**d**, Representative DIC images of *E. coli* MG1655 cells transformed with the indicated constructs grown for 4 h either in the dark or under 460 nm light (5 W/m^2^) illumination. **c**,**e**,**f**, Quantification of cell length distribution for the indicated samples and conditions. NC, pBLADE-empty. **g**, Representative DIC images of *E. coli* KC717 cells transformed with the indicated constructs at the indicated time points. Induction indicates 460 nm light (5 W/m^2^) for the cells transformed with pBLADE-RodZ and 0.2% arabinose for the cells transformed with an empty pBAD33 from which the P_BAD_ promoter was removed (pBAD#). Recovery indicates darkness for the cells transformed with pBLADE-RodZ and growth in a medium without arabinose for the cells transformed with pBAD#. **h**, Quantification of the cell roundness for the samples and conditions in (**g**). **b,d,g**, Scale bar, 5 mm. **c**,**e**,**f**,**g**, Values represent mean ± s.d. of *n=3* independent experiments. **b**-**h**, BLADE variant: FP4 driven by the J23101** promoter.

### Characterization of *E. coli* genes with unknown function in terms of intracellular localization and effect on growth and morphology

The *E. coli* genome contains 4623 genes, 35% of which currently lack experimental evidence of function^36^. Since light is particularly well-suited for medium to high-throughput studies due to its low cost, scalability, and effortless application, we used BLADE to characterize some of these genes in terms of their intracellular localization and effect on cell growth and morphology. We randomly selected 34 completely uncharacterized genes and included 5 additional genes, for which some information was available: *ydaT*, which was shown to lead to cell elongation and reduced survival when overexpressed in *E. coli*^37^; *ydiY*, which was shown to be induced by acid and was predicted to be an outer membrane protein^38^; *ycbK* (renamed MepK), which was shown to be a murein hydrolase involved in cell wall synthesis^39^; *yehS*, whose downregulation has been shown to improve the growth of *E. coli* in n-butanol and n-hexane^40^; and *yebE*, which was shown to be induced by copper in a CpxA/CpxR-dependent manner^41^, and was predicted to be localized to the inner membrane^42^. Importantly, fluorescence microscopy-based localization studies have not been so far carried out for any of these 39 genes. Since fusion to a fluorescent protein (FP) could alter or impair the function of the gene products, we cloned each of the 39 genes in native form into pBLADE (Fig. 4a). However, in order to monitor the localization of the gene products in *E. coli* cells, we additionally cloned for each gene N- and C-terminal fusions to sfGFP (Fig. 4a). We tested both termini because it is known that the terminus at which the FP is fused often plays a role in determining whether the fusion protein maintains the same localization as the native one^43–45^. In total, we constructed a library of 117 plasmids. Those bearing the native genes were subjected to growth assays and differential interference contrast (DIC) microscopy, while those bearing the fusions to sfGFP were subjected to fluorescence microscopy (Fig. 4a). Before performing experiments, we applied bioinformatics and computational structural biology approaches to predict the function and localization of the 39 selected genes (Fig. 4a). We used three different tools (Argot2.5^46^, PANNZER2^47^, and DeepGoPlus^48^) that predict protein function and localization from amino acid sequence information only and one protein 3D modeling tool (Phyre2^49^) that uses this information as well as secondary structure prediction to find a template structure that best represents the submitted protein for 3D modeling (Supplementary Table 1). We generated a consensus table for localization and function taking the predictions shared by at least two out of the four methods (Supplementary Table 2). A consensus was found for 14 out of 39 genes for functional prediction and for 21 out of 39 for localization prediction. We first analyzed the effect of overexpressing the native proteins on bacterial growth. We found six genes whose products significantly affected the growth of MG1655 cells: three positively (*yahC*, *yebE* and *yebY*) and three negatively (*yhhM*, *yjeO* and *ypaB*; Fig. 4b and Supplementary Fig. 7a). Interestingly, *yebY* is predicted to have transaminase activity (Supplementary Table 2), which could explain why cells overexpressing it grow faster. To assess if any of the 39 genes caused morphological changes, we performed DIC microscopy on MG1655 cells exposed to light for 4 hours. While most genes did not cause morphological alterations, two led to cell elongation (*ydaT* and *ydhL*) and one to cell lysis (*yhcF*; Fig. 4c). Our results thus confirm previous observations on the effect of *ydaT* overexpression on cell morphology^37^, and further indicate that *ydhL* may be involved in cell division. Since *ydcF* overexpression caused cell death in this assay, we additionally measured the OD_600_ of the cell culture after 4 hours of growth in the incubator and found that it was indeed reduced compared to that of the cultures overexpressing *ydaT* and *ydhL* as well as compared to cells transformed with empty pBLADE (Supplementary Fig. 7b). Notably, *yhcF* did not cause growth defects in the assay performed in the 96-well plate. However, it is known that bacteria grow slower in a 96-well plate than in a flask, due to the lower oxygen exchange and shaking. Therefore, it cannot be excluded that the 6 genes found to affect growth in the 96 well-plate assay form only a partial list, and that other genes among the selected 39 may also affect *E. coli* growth when overexpressed.

**Fig. 4.**
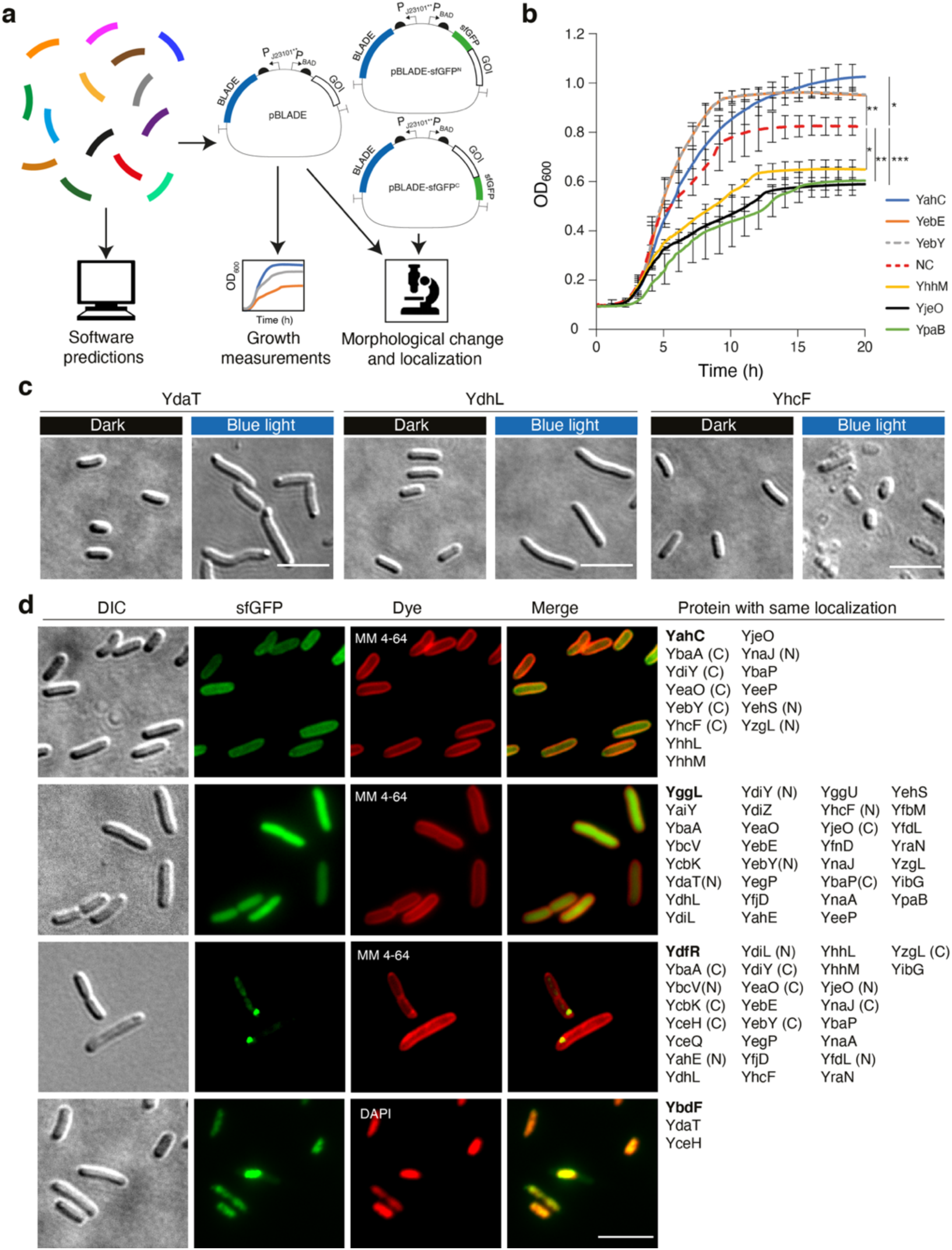
**BLADE facilitates the characterization of *E. coli* genes with unknown or poorly defined function. a**, Overview of the workflow. pBLADE-sfGFP^N^, plasmid for sfGFP N-terminal fusion; pBLADE-sfGFP^C^, plasmid for sfGFP C-terminal fusion. **b**, Growth curves of *E. coli* MG1655 cells transformed with pBLADE carrying the indicated proteins. NC, cells transformed with the empty plasmid. Values represent mean ± s.d. of *n=3* independent experiments. Single asterisk (*), p-value<0.5 (two-tailed, homoscedastic Student’s *t* test); double asterisk (**), p-value<0.01 (two-tailed, homoscedastic Student’s *t* test); triple asterisk (***), p-value<0.001 (two-tailed, homoscedastic Student’s *t* test). **c**, Representative DIC images of *E. coli* MG1655 cells transformed with pBLADE carrying the indicated proteins grown for 4h either in the dark or under 460 nm light (5 W/m^2^) illumination. **d**, Representative images of *E. coli* MG1655 cells transformed with pBLADE-sfGFP^N^ carrying the proteins indicated in bold in the right-most column grown for 4h under 460 nm light (5 W/m^2^) illumination. (N), localization obtained only with N-terminal fusion; (C), localization obtained only with C-terminal fusion. The membrane was stained with the MM 4-64 dye and the nucleoid with DAPI. **c**,**d**, Scale bar, 5 μm.

To study the localization of the uncharacterized genes, we performed fluorescence microscopy. As expected, not all gene products tolerated fusions to either terminus (Supplementary Table 2). For some, fluorescence was barely detectable for one of the two fusions, and for others the localization was not the same for both fusions (Supplementary Fig. 8a, b and Supplementary Table 2). All in all, we found 3 genes whose products co-localized with the nucleoid, 14 that co-localized to the cytoplasmic membrane, and 26 that formed foci (Fig. 4d). While the localization alone is not sufficient to reveal the function of the 39 genes, it gives important information and, for some of the genes, suggests a potential mechanism of action. For instance, *ydaT*, which is reported to be a toxin^37^, may exert this function by binding and inhibiting DNA gyrase, since we found it co-localized on the nucleoid. Other toxins that inhibit DNA gyrase and co-localize to the nucleoid have been described^50, 51^.

### The mechanism of BLADE-mediated blue light-inducible gene expression involves the formation of intracellular protein aggregates in the dark

Wild type AraC and BLADE are substantially different in their mode of action: AraC is always a dimer that, in the absence of arabinose, binds the I_1_ and O_2_ half-sites and, in the presence of the sugar, binds the I_1_ and I_2_ half-sites (Fig. 1a). In contrast, BLADE is monomeric in the dark and dimeric under blue light illumination (Fig. 5a). It is hard to predict the 3D structure of the light-induced BLADE dimer. In principle, dimeric BLADE could assume a conformation resembling either that of the AraC dimer free of arabinose, or that of the arabinose-bound dimer, or even a different conformation not found in the natural protein that would nonetheless favor interaction with I_1_ and I_2_. All the data we obtained strongly suggest that dimeric BLADE preferentially assumes a conformation that leads to its interaction with the I_1_ and I_2_ half-sites. In particular, the increase in reporter gene expression after illumination can be explained only if BLADE contacts the I_2_ half-site and recruits the RNA polymerase at the P_BAD_ promoter. The I_1_ half-site is likely contacted also by monomeric BLADE, given that a single DBD of AraC was shown to bind to it *in vitro*^12^. However, this is not sufficient for recruiting the RNA polymerase to the P_BAD_ promoter^52^. To further prove that the I_2_ half-site is contacted *in vivo* by BLADE in the presence of blue light, we constructed a modified pBLADE plasmid, in which the I_2_ half-site was cloned in inverse orientation, while keeping the -35 region of the P_BAD_ promoter untouched (Supplementary Fig. 9a). In this case, there was no significant difference in mCherry levels between dark and light samples (Supplementary Fig. 9b).

**Fig. 5.**
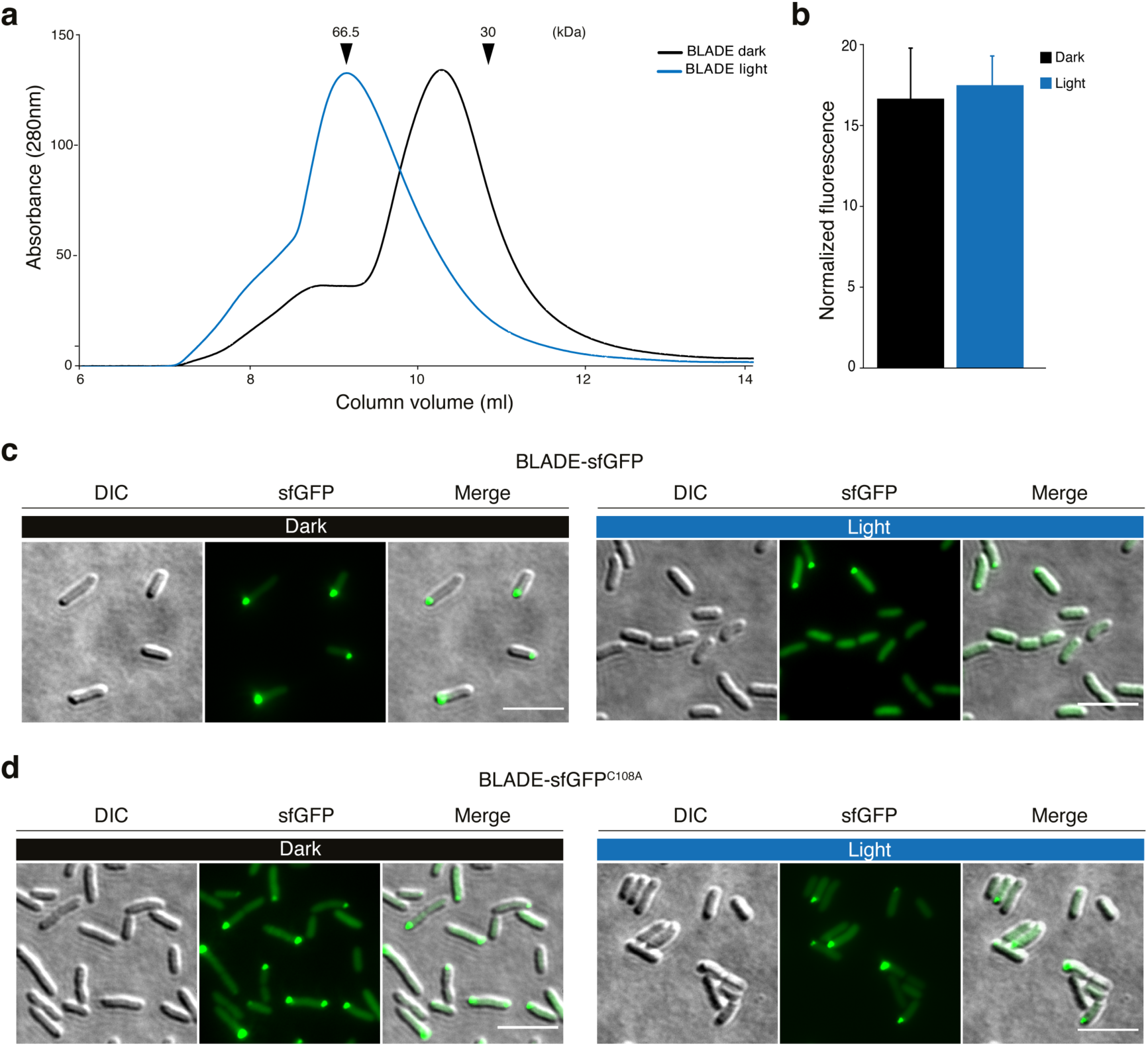
**BLADE-mediated light-induced gene expression involves the formation of aggregates in the dark and of dimers in the light. a**, SEC performed with purified BLADE in the dark or illuminated with 460 nm light (5 W/m^2^) for 30 minutes at 4°C. **b**, GFP fluorescence intensity measured in *E. coli* MG1655 cells transformed with a modified pBLADE in which BLADE was C-terminally fused with sfGFP grown for 4h in the dark or under 460 nm light (5 W/m^2^) light. **c**,**d**, Representative microscopy images of *E. coli* MG1655 cells expressing the indicated BLADE variant C-terminally fused to sfGFP grown for 4h in the dark or under 460 nm light (5 W/m^2^) light. Scale bar, 5 μm.

In *Neurospora crassa*, the organism in which it is naturally expressed, VVD is actually degraded in the dark^53–55^. We asked whether VVD may trigger the degradation of BLADE in *E. coli* cells in the dark, which could add another layer of regulation to the system and contribute to its tightness. To address this question, we fused sfGFP to the C-terminus of BLADE to avoid any interference with dimerization and performed flow cytometry to measure BLADE levels in cells kept in the dark and exposed to blue light for 4 hours. The sfGFP levels were comparable in both conditions (Fig. 5b). However, fluorescence microscopy revealed the presence of bright fluorescent foci in half of the cells kept in the dark (Fig. 5c and Supplementary Fig. 9c), while less than 20% of the illuminated cells showed foci (Fig. 5d and Supplementary Fig. 9c). To investigate the nature of the foci, we performed fluorescence recovery after photobleaching (FRAP) experiments and found the foci to be static (Supplementary Fig. 9d), suggesting they are aggregates rather than functional liquid droplets. It has been previously shown that VVD transitions between locally unfolded and folded states and that light shifts the half-life of the transition from about 5 minutes to 6 hours^56^. It was suggested that simultaneous unfolding of several structural elements of VVD could lead to aggregation in the dark^56^. The aggregates we observed in *E. coli* could, therefore, be due to the VVD moiety in BLADE. To prove that the aggregates are related to the light response of VVD, we mutated the adduct-forming cysteine to alanine (VVD^C108A^) within BLADE. We expected the mutant to show aggregates also under blue light illumination, since VVD^C108A^ is not responsive to light. Indeed 40% of the cells presented aggregates both in the dark and when illuminated with blue light for 4 hours (Fig. 5e and Supplementary Fig. 9c).

Finally, to investigate whether light leads to the dispersion of previously formed foci, we performed time-lapse fluorescence microscopy to follow individual foci over time in illuminated cells. If light actively disperses the aggregates, foci in individual cells should disappear. Alternatively, the aggregates may remain intact under blue light illumination, but form less frequently in newborn cells. We found that the aggregates do not disperse, but are instead asymmetrically segregated during cell division (Supplementary Video 2). Newborn cells contain either no foci or foci much smaller than those found in cells kept in the dark (Supplementary Video 2).

### Expanding the family of BLADE TFs

In principle, BLADE could have been designed using other light-inducible dimerization domains. Moreover, the position of this domain with respect to the DBD of AraC may not need to reflect that found in the wild type protein. To test if other functional combinations with different characteristics could be identified, we generated a much larger set of samples for characterization, with a library size significantly larger than the one described earlier. As a light-inducible dimerization unit, we included not only VVD, but also the Light Oxygen Voltage (LOV) domain of *Vaucheria frigida* Aureochrome1 (VfAu1)^57, 58^, which is naturally found C-terminally to a bZip DBD^58^, and which, like VVD, homodimerizes upon blue light stimulation^59, 60^. To assess the functionality of the chimeric transcription factors (cTFs), we used only the P_BAD_ promoter (I_1_-I_2_ half-sites) and removed the upstream regulatory elements (O_1_ and O_2_ half-sites)^61^. Since the results with the initial VVD-AraC fusions showed that the strength of the constitutive promoter driving their expression played an important role in determining the light/dark fold change (Fig. 1f), we systematically explored how the expression levels of the cTF affected mCherry levels in the dark and after blue light illumination. To this aim, we first used an isopropyl-β-D-thiogalactopyranoside (IPTG)-inducible promoter^62^ to achieve various levels of expression of the cTF, with the goal of finding the most appropriate expression level. This identification of appropriate or ‘optimal’ cTF expression levels to achieve a certain output (e.g. highest fold change, or certain levels of dark or light-induced expression) is the first step in our 2-step method (Fig. 6a). For this, an IPTG-inducible promoter is used to cover a wide range of cTF concentrations—a step that only requires a single genetic construct. The second step maps the transcriptional strength of the IPTG induction levels to constitutive promoters. This step only needs to be performed once for a given inducible promoter. Using constitutive promoters allows for future uses of the optimized systems and eliminates the need for inducer molecules. We employed the IPTG-inducible promoter as part of single-plasmid systems that can be assembled in one-pot Golden Gate cloning reactions comprised of easily adaptable components. This allows for the characterization of these and other cTFs by enabling the exchange of every functional genetic component (Supplementary Fig. 10). In principle, one may expect that different possible scenarios could arise from the influence of the cTF concentration on the output expression (Supplementary Fig. 11). For example, Scenario 1 represents the case in which the higher the cTF concentration, the higher the output will be, both in the dark and after illumination, maintaining the fold change relatively constant. Scenario 2 represents the case in which a concentration threshold exists, after which there is a reduction in the light-induced fold change of the output. This effect could be due, for example, to resource limitations in the cells that express the cTF and the output gene. Scenario 3 corresponds to a different effect—one that also implies the existence of optimal intermediate cTF concentrations. Here, high cTF concentrations do not alter the output expression in the light, but they instead cause the dark state to increase, for instance due to the formation of dimers in the dark. We used a wide range of inducer concentrations to capture these potential scenarios, and focused on the output light/dark fold change, an important feature of light-inducible proteins. Depending on the application, other properties such as high output expression or low dark state might be more relevant. To characterize many individual samples, under the same light input conditions, a novel high-throughput light induction device was needed. We therefore developed a light induction device which can be used for standard 96-well microtiter plates in which the light input for every well can be steered individually (Fig. 6b). The setup comprises a custom-made printed circuit board (PCB) with 96 individual light emitting diodes (LEDs) of three different wavelengths (red, green and blue). Each LED can be controlled individually using a microcontroller, enabling the exposure of each well to the same light intensity, a crucial aspect for the characterization of the cTFs. A milled metal plate placed in between the PCB and the LEDs dissipates the heat produced by the light induction device. A 3D-printed microplate adapter on top of the metal plate allows for the precise positioning of the 96-well plate. The high-throughput characterization allowed us to calculate IPTG dose-response curves of the same construct receiving the same light input as well as grown in the dark (Fig. 6a). In addition, it allowed us to test the N- and C-terminal positioning of the light-inducible dimerization unit, as well as different linkers connecting the two domains, which would not have been feasible without this technical setup (Fig. 6c).

**Fig. 6.**
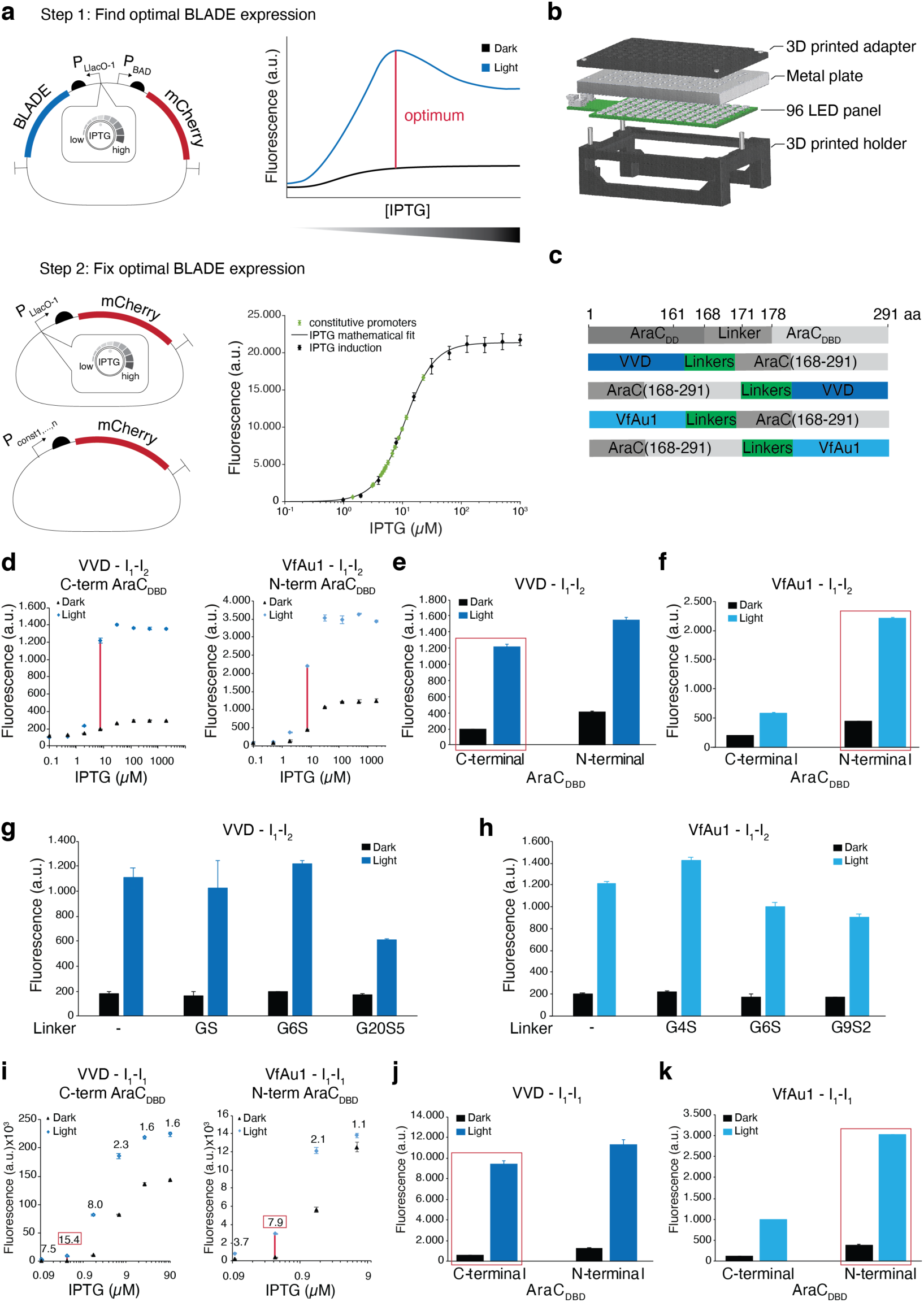
**Engineering an optimized and expanded family of BLADE TFs. a**, A two-step protocol was used to optimize the expression levels of the TFs. Upper panel: left, the plasmid used in step 1 contains IPTG-inducible chimeric BLADE transcription factors (cTFs) and an mCherry reporter under the P_BAD_ promoter. Upstream regulatory sequences (O_1_ and O_2_ half-sites) have been deleted. Each IPTG concentration induces a different BLADE cTF level with a corresponding light/dark mCherry expression profile. Measuring these profiles at varying IPTG concentrations allows for the identification of a profile that is optimal with respect to a desired property. Right, example of a scenario for which the highest mCherry levels are obtained at intermediate cTF levels (red line). Bottom panel: In step 2, the range of transcriptional rates induced by different IPTG concentrations is mapped to the rates of constitutive promoters in a library (right), allowing for the identification of a constitutive promoter that matches the desired optimal BLADE expression level. **b**, Light induction setup for 96-well microtiter plates containing a panel of 96 light emitting diodes (LEDs), 3D-printed holders and a metal plate for heat dissipation. **c**, Domain composition of the engineered light-inducible dimerization domain (VVD and VfAu1)-AraC_DBD_ fusion constructs. **d**, Examples of IPTG dose-response curves obtained with BW25113 *ΔaraC +lacYA177C* cells transformed with the indicated constructs (VVD::G_6_S::AraC_DBD_ and AraC_DBD_::(G_4_S)_5_::VfAu1). The highest light/dark fold change is indicated with a red line between the corresponding data points. **e**, mCherry fluorescence intensity in BW25113 *1araC +lacYA177C* cells transformed with the VVD::G_6_S::AraC_DBD_ fusion (N-terminal) and the AraC_DBD_::G_4_S::VVD fusion (C-terminal) in the presence of 7.81 µM IPTG. The samples in the red box are taken from the dose response curve shown in (**d**). The IPTG dose-response curve of AraC_DBD_::G_4_S::VVD is shown in Supplementary Fig. 14. **f**, mCherry fluorescence intensity in BW25113 *1araC +lacYA177C* cells transformed with the VfAu1::(G_4_S)_5_::AraC_DBD_ fusion (N-terminal) and the AraC_DBD_::(G_4_S)_5_::VfAu1 fusion (C-terminal) grown in the presence of 7.81 µM IPTG. The samples in the red box are taken from the dose response curve shown in (**d**). The IPTG dose-response curve of VfAu1::(G_4_S)_5_::AraC_DBD_ is shown in Supplementary Fig. 15. **g**, mCherry fluorescence intensity in BW25113 *1araC +lacYA177C* cells transformed with a small library of C-terminal AraC_DBD_ fusions with VVD with the indicated linkers between the two domains. The corresponding IPTG dose-response curves are shown in Supplementary Fig. 14. **h**, mCherry fluorescence intensity in BW25113 *ΔaraC +lacYA177C* cells transformed with a small library of C-terminal AraC_DBD_ fusions with VfAu1 with the indicated linkers between the two domains. The corresponding IPTG dose-response curves are shown in Supplementary Fig. 15. **i**, Same as in (**d**) but with a synthetic P_BAD_ promoter containing two copies of the I_1_ half-site. The highest light/dark fold change is indicated with a red line between the corresponding data points. **j**, Same as in (**e**) but with a synthetic P_BAD_ promoter containing two copies of the I_1_ half-site and 0.391 µM IPTG. The IPTG dose-response curve of AraC_DBD_::G_4_S::VVD is shown in Supplementary Fig. 16. **k**, Same as in (**f**) but with a synthetic P_BAD_ promoter containing two copies of the I_1_ half-site and 0.391 µM IPTG. The samples in the red box are taken from the dose response curve shown in **(i**). The IPTG dose-response curve of VfAu1::(G_4_S)_5_::AraC_DBD_ is shown in Supplementary Fig. 17. All values represent mean ± s.d. of *n=2 or 3* independent experiments. In all experiments, cells were grown for 5h either in the dark or under 460 nm light illumination.

For this characterization we used *E. coli* strain BW25113 *ΔaraC* from the KEIO collection^63^, and integrated *lacYA177C* into the *attB* site for facilitated IPTG diffusion^30^. We first tested the constructs in the library with variable order between AraC_DBD_ and the light-inducible dimerization unit, but constant linker. Our 96-well light induction setup allowed us to use a wide range of IPTG concentrations, from no induction to a concentration of 2 mM IPTG. For all constructs, the highest fold change was reached at intermediate mCherry expression levels (indicated with red lines in two examples shown in Fig. 6d). Placing AraC_DBD_ at the C-terminus led to higher fold changes for VVD-based constructs, mainly due to lower mCherry expression in the dark (Fig. 6e). For VfAu1, the opposite was true (Fig. 6f). Next, we investigated the effect of linker length on the cTFs. Based on the results obtained with the first library, we placed AraC_DBD_ C-terminally for the VVD-based constructs, and N-terminally for those based on VfAu1. We selected a set of linkers from a previous report^64^. Additionally, we cloned a linker-free variant for each of the dimerization domains, where the domains where directly fused with each other. We found that linker lengths from zero to 7 amino acids gave rise to the highest fold change for both photosensors (Fig. 6g, h). All these functional fusions expand the family of BLADE TFs.

Since in future biological applications BLADE should be constitutively expressed to dispense of the use of any chemical inducer, as was the case in our initial experiments, we aimed to find constitutive promoters that would give rise to expression levels comparable to those obtained with various IPTG concentrations. We therefore cloned in the same plasmid bearing the IPTG-inducible promoter a library of constitutive promoters^65^, as well as an additional weaker constitutive promoter variant to cover very low expression levels. To minimize the potential influence of individual promoters on mRNA transcription and translation initiation, we used a ribosome binding site (RBS) containing an insulating ribozyme (RiboJ)^66^. Plotting the mCherry fluorescence levels obtained with the constitutive promoters and with the IPTG-inducible promoter at different IPTG concentrations in the same plot, it is possible to find the constitutive promoter that best matches the expression from the IPTG-inducible one at the desired IPTG concentration (Fig. 6a). While the characterization was performed in an *E. coli* strain in which the arabinose operon was deleted, the results do not change if a strain with endogenous *araC* is used (Supplementary Fig. 12).

### A synthetic P_BAD_ promoter comprising two I_1_ half-sites is light-inducible and achieves higher light/dark fold changes

We performed the same systematic characterization of BLADE family members using a synthetic P_BAD_ promoter, where the weak affinity I_2_ half-site was exchanged with a second copy of the high affinity I_1_ half-site (Supplementary Fig. 13). This synthetic promoter is insensitive to arabinose, as it is constitutively active when used with wild type AraC^67^. We asked whether it could be made light-inducible instead. Results with this promoter were consistent with those obtained with the synthetic P_BAD_ promoter consisting only of the I_1_ and I_2_ half-sites (Fig. 6j, k), confirming our findings regarding the position of the domains and the linkers within the cTFs. Interestingly, the maximal dark/light fold change for the same cTFs was higher compared to that obtained with the I_1_-I_2_ synthetic promoter. High IPTG concentrations led to toxic amounts of mCherry expression and were, therefore, indistinguishable for dark and light induction in most cases. Careful adjustment of the cTF concentration is required to achieve the desired light inducibility, which then unlocks an expression system with an even higher expression level and fold change than the one based on the original I_1_ and I_2_ half-sites.

## Discussion

AraC is among the best studied bacterial transcriptional regulators, and the P_BAD_ promoter is one of the inducible promoters most often employed in microbiology and synthetic biology. We have developed an entire family of AraC-derived TFs, which we call BLADE, that activate transcription from the P_BAD_ promoter in response to blue light instead of arabinose. BLADE TFs are compatible with previously constructed strains carrying the P_BAD_ promoter at an endogenous locus to drive the expression of a gene of interest, allowing microbiologists to readily perform optogenetic experiments without the need to construct anything new – transformation of the strain with pBLADE is the only requirement. Moreover, since we constructed pBLADE using pBAD33 as template (Supplementary Fig. 2a), microbiologists who wish to implement optogenetic control of their gene of interest can simply re-clone it into pBLADE using the same restriction enzymes previously employed with pBAD33. Another advantage is that the resistance cassette and origin of replication of pBLADE are identical to those of pBAD33, thus ensuring compatibility with other previously constructed plasmids that should be co-transformed with pBLADE. We additionally envisage that a plasmid carrying BLADE under a constitutive promoter may be combined with previously constructed arabinose-inducible plasmids as long as the origin of replications and resistances are compatible. This strategy would require no cloning and guarantee full compatibility with established plasmids.

While many other light-inducible TFs have been developed to date^61, 68–72^, some of which featuring extremely high dark/light fold changes^61, 72^, we explicitly aimed to engineer a system based on a well-known and pervasive TF (namely AraC) that is particularly suited for microbiological applications, thus stimulating the use of optogenetics in microbiology. We took special care to engineer BLADE with minimal leakiness. Often, leakiness has been assessed by comparing the levels of reporter expression in the dark to those in the light. However, this does not take into account whether the expression in the dark state is already too high compared to the expression in the absence of the TF. Therefore, in the case of BLADE, minimal leakiness was demonstrated by comparing its activity in the dark with expression obtained with the same plasmid deprived of the TF (Fig.1f-i and Supplementary Fig. 2b). We put BLADE to the test by expressing several functional *E. coli* proteins whose overexpression causes morphological changes to the cells and showed that, in the dark, cells are indistinguishable from the control (Fig. 3).

Light can be easily switched on and off. We showed that BLADE allows controlling a phenotype in a fully reversible manner, using rod shape as an example (Fig. 3g, h). In contrast to the fast reversibility achievable with pBLADE, the chemical inducer arabinose, even after being washed off, remained inside the cells, committing them to become even more rod-shaped instead of going back to the spherical morphology. Moreover, light could be locally applied to create mixed populations to directly compared the effects of morphology on the fitness of cells in different environments.

Another key feature of BLADE is that its leads to a homogenous response in a cell population (Supplementary Fig. 3), in contrast to the heterogenous activation of P_BAD_^73^. While heterogeneity can be overcome by overexpression of either the arabinose transporter AraE^74^ or LacY^75^, a transporter with relaxed specificity, the usage of the promoter is then limited to these engineered strains or requires co-transformation with a plasmid encoding the transporter.

To showcase the utility of light induction in medium to high-throughput studies, we used BLADE to overexpress 39 genes randomly selected from those with unknown or poorly defined function. We applied four bioinformatic tools to predict their function and localization. While it was possible to find a consensus prediction for 53% of the genes in case of localization, only for 35% of the genes was a consensus found for function prediction. Even in this case, the prediction remained relatively vague (ligase, transporter, DNA binder, etc.; see Supplementary Table 1). This highlights how computer-based predictions cannot yet replace detailed biochemical characterization, which remains essential to assign a function to a protein.

Previous reports on light-inducible TFs based on VVD have employed untagged versions, since the scope was to quantify the reporter gene expression output^5, 6, 14, 72^. We also did not include any FP in the BLADE construct at the beginning, since visualization of BLADE localization was not important and the fusion may have not been as active as the untagged version. It was, thus, surprising to see that BLADE is not simply cytoplasmic as it may be expected (Fig. 5c). Formation of aggregates in the dark is in good accordance with previous studies^56, 76^. While we did not analyze VVD alone, we speculate that the aggregates reflect a property of VVD, not AraC_DBD_. Evidence in this respect comes from the results with BLADE bearing a mutated VVD (VVD^C108A^), which is insensitive to light and forms aggregates also under blue light illumination (Fig. 5d). While in *N. crassa* VVD is degraded in the dark, in *E. coli* the protein forms aggregates, which may effectively have an impact similar to degradation in its rendering the protein inactive. The advantage of inactivation by sequestration rather than degradation is that the protein can be quickly released from the aggregates and activated when needed, without the delay that would result from a novel round of gene expression. To date, many proteins have been shown to localize to intracellular bodies, however, often it is not clear if they constitute functional entities, such as liquid droplets, storage bodies or aggregates^77^. For the protein stored in the aggregates to be quickly activated, its association with these aggregates should be dynamic. When followed over time in time-lapse microscopy, we found that the aggregates formed by BLADE in the dark did not disperse in the same cell once it was illuminated with blue light (Supplementary Video 2). The aggregates were polarly localized and asymmetrically segregated to only one of the daughter cells (Supplementary Video 2). This is in line with a previous report on asymmetric segregation of protein aggregates in *E. coli*^78^. These data suggest that the foci formed by BLADE are indeed dysfunctional aggregates and that, when cells are illuminated, the probability of their *de novo* formation strongly decreases. Notably, the aggregates cannot be the only mechanism in place that controls the activation of the P_BAD_ promoter by BLADE, since we found them only in about half of the cells (Supplementary Fig. 9c). We believe they add a layer of regulation to the system, contributing to its tightness. However, the mechanism of BLADE-mediated gene expression involves dimer formation and consequent occupancy of the I_2_ half-site, which recruits the RNA polymerase.

When engineering new TFs, we found that not only is an appropriate protein engineering approach necessary, but also that the adjustment of the TF concentration is critical if we wish to achieve optimal functionality. While this is intuitive for low TF concentrations that might be insufficient to generate a biological response, we also found that, if concentrations exceed certain levels, the functionality of the TF may deteriorate (Fig. 6d) or disappear altogether (Fig. 6i). For the selected metric of light/dark fold change, intermediate TF expression levels always led to the highest, and hence the most optimal, values. Our high-throughput approach and novel 96-well light induction device made this possible. By calibrating the IPTG dose-response of an inducible promoter to a large set of constitutive promoters, the desired TF expression levels can be fixed, dispensing the need for an inducer (Fig. 6a). This mapping is essential, because it dramatically reduces the experimental demand for cloning of constructs, as only one construct needs to be cloned per TF. While this approach holds great promise for the optimization and development optogenetic systems, as demonstrated here, we expect that it will also be useful for the development of other transcriptional regulators as well as for systematically bringing the expression levels of various components in biological circuits to their optimal levels.

In this article we have demonstrated several features enabled by light induction, including reversibility, which opens up the possibility of using dynamic inputs for probing biological phenomena. In particular, pulsatile inputs that alternate between dark (OFF) and maximum intensity (fully ON) can be used to achieve effects that cannot be realized with graded intensity light. For example, it has been shown that pulsatile inputs lead to reduced cell-to-cell variability in gene expression^79^. In fact, by adjusting the duty-cycle (defined as the fraction of the time that the light is fully ON), one can even tune the amount of cell-to-cell variability, providing a new control modality for exploring cell-to-cell variability and stochastic gene expression. This type of pulsatile input was also recently shown to enhance the biosynthesis of products in engineered cells, enabling a new manner of bioreactor operation in which enzyme expression is tuned to increase fermentation yield^80^.

Taken together, the features of BLADE, its ease of adoption and usage, and its low cost should bring the many benefits of optogenetic manipulation to the field of microbiology, enabling new and exciting discoveries.

## Methods

### Strains, media and reagents

The strains used in this study are listed in Supplementary Table 3. For experiments shown in Figures 1, 3-5, and Supplementary Figures 3, 4, 5 and 8, the cultures were grown in autoclaved Tryptone Broth (TB; 10 g l^-1^ Tryptone, 5 g l^-1^ NaCl, 1 mM NaOH). For the bacterial photography experiments shown in Figure 2, and Supplementary Figure 7, the cultures were grown in autoclaved LB-Miller medium. For experiments shown in Figure 6 and Supplementary Figures 12, 14-17, the cultures were grown in autoclaved LB-Miller medium for strain propagation and in sterile-filtered M9 medium supplemented with 0.2% casamino acids, 0.4% glucose, 0.001% thiamine, 0.00006% ferric citrate, 0.1 mM calcium chloride, 1 mM magnesium sulfate for all gene expression experiments. In experiments in which the plasmid had to be maintained, the medium was supplemented with 34 μg ml^-1^ chloramphenicol (Sigma-Aldrich Chemie GmbH). IPTG, rifampicin and doxycycline were purchased from Sigma-Aldrich Chemie GmbH.

### Constructions of strains and plasmids

To integrate *lacYA177C* into the *attB* site of BW25113^63, 81^ we used *λ* integrase expressed from pJW27 using plasmid pSKA27 containing *lacYA177C*, FRT-flanked *kanR* from pKD13 ligated into XbaI-cut pFL503^82^ and a sequence identical to the genome regions for *attB* integration. pSKA27 was cut with NotI, and the 4229 bp band gel was purified and circularized before transformation into pJW27-containing cells. For integration, pJW27 was transformed into *E. coli* BW25113 and selected at 30°C on LB-Agar plates containing chloramphenicol for expression of *λ* integrase. A single colony was used to inoculate 5 ml of LB broth containing chloramphenicol, and the culture was grown at 30°C in a water bath with shaking. The cells were then moved to 42°C for 15 min, before incubating on ice for 15 min. Cells were transformed with the integration construct using the previously described transformation protocol. The fusion VVD-AraC proteins FP1-5 were first subcloned into pDK12^83^ using the NcoI and NotI restriction sites. The *vvd* gene carrying the N56K and C71V mutations and coding for a VVD protein missing the first 36 amino acids was PCR-amplified out of plasmid pGAVPO (gift from Yi Yang; East China University of Science and Technology). The *araC* fragments were amplified from pBAD33. To clone the fusions, a two-step protocol was followed. In the first step, the two parts were separately PCR-amplified. After the purification of the PCR products, the two fragments were fused together in the second PCR step, and then cloned into pDK12 with NcoI and NotI restriction enzymes, yielding plasmids pDK12(FP1-5). Next, the DNA sequences coding for the fusion proteins FP1-FP5 were PCR-amplified using primers 15 and 16 and cloned into pBAD33 (previously deprived of AraC via PCR using primers 13 and 14, yielding the negative control plasmid pBLADE-empty) linearized with ClaI. The J23101 promoter was included in the forward primer. The obtained pBAD33-derived plasmids are called pBLADE(FP1-5). The *mCherry* gene codon-optimized for expression in *E. coli* was synthesized (IDT) and cloned into pBLADE with SacI and HindIII restriction enzymes. We subsequently generated pBLADE(FP7)-mCherry by inserting an additional GS linker between VVD and AraC with a site-directed mutagenesis protocol using pBLADE(FP4)-mCherry as template with primers 17 and 18. The primers, designed with the QuikChange Primer Design website, were used to amplify the plasmid. The PCR reactions contained 8% of DMSO to allow proper annealing of the primers to the template DNA. Similarly, pBLADE(FP6)-mCherry was generated by removal of the GS linker from the same template with the same protocol previously described, using primers 19 and 20. To construct the positive control with full-length AraC, pBLADE was linearized with ClaI, the *araC* gene was PCR-amplified with primers 21 and 22 and then cloned into the backbone, yielding pBLADE(AraC_WT_)-mCherry. The mutations and deletions leading to promoters J23101* and J23101** (Supplementary Figure 1) generated spontaneously during growth of bacterial cultures transformed with pBLADE(FP4)- mCherry. These promoters have been subsequently cloned in all other pBLADE plasmids by PCR-amplification with primers 23 and 24. The backbones were PCR amplified with primers 25 and 26, yielding pBLADE(FP1*/FP7*)-mCherry and pBLADE(FP1**/FP7**)-mCherry. The promoters were cloned into pBLADE(AraC_WT_)-mCherry by overlapping PCR starting from pBLADE(AraC_WT_)- mCherry as template with primers 27 and 28, yielding pBLADE(AraC_WT_*/AraC_WT_**)- mCherry. The DNA sequence coding for eYFP-MinD was PCR-amplified out of pSR-4^84^ with primers 31 and 32 and cloned into pBLADE(FP6*) via Gibson Assembly after having amplified the backbone (pBLADE(FP6*)-mCherry) with primers 33 and 34. The *sfgfp* gene was PCR-amplified with primers 35 and 36 from plasmid pHR-scFv-GCN4-sfGFP-GB1-NLS-dwPRE (gift from Ron Vale; Addgene plasmid # 60906; http://n2t.net/addgene:60906; RRID:Addgene_60906) and cloned in pBLADE(FP6*)-mCherry via Gibson Assembly after amplification of the backbone with primers 37 and 38, yielding pBLADE(FP6*)-sfGFP. The DNA sequence coding for MinD^Δ10^ was amplified out of pBDV-13^18^ with primers 39 and 40 and cloned via Gibson Assembly into pBLADE(FP4**)-mCherry previously amplified with primers 41 and 42. The *rodZ* and *mreB* genes were PCR-amplified from genomic DNA isolated from *E. coli* MG1655 using primers 43 and 44 and 45 and 46 respectively. Cloning into pBLADE(FP4**) was achieved via Gibson Assembly after amplification of the plasmid backbone with primers 33 and 34. pBAD# (pBAD33 deprived of the P_BAD_ promoter and mCherry) was cloned via Gibson Assembly by amplification of pBLADE(FP6**)-mCherry with primers 47 and 48, digestion of the linearized plasmid with SacI and following ligation. The 39 genes with unknown or poorly defined function were PCR-amplified from genomic DNA isolated from *E. coli* MG1655 using the primer pairs listed in Supplementary Table 5. The backbone pBLADE(FP6**) was amplified with primers 33 and 34 to insert the first 16 genes, and with primers 49 and 50 to insert the others. These primers allow maintaining start and stop codon on the plasmid backbone. To create the library with the fusion to sfGFP, the first 16 genes in the list in Supplementary Table 1 were amplified with primers that included a GS linker at their N or C-terminus, and cloned in the backbone pBLADE(FP6**)-sfGFP amplified with primers 33 and 51 (N-terminal fusions) and 52 and 34 (C-terminal fusions). For the remaining genes, primers 50 and 54 (N-terminal fusions) and 49 and 53 (C-terminal fusions) were used. For protein purification, the BLADE FP6 construct was PCR-amplified with primers 55 and 56 from pBLADE(FP6*)-mCherry and cloned into pET28a with BamHI and NotI restriction enzymes, yielding to pET28a-FP6. In order to invert the orientation of the I_2_ half-site, the plasmid pBLADE(FP6*)-mCherry was amplified with overlapping PCR with primers 57 and 58 generating the plasmid pBLADE_I_2_rev_(FP6*)-mCherry. Note that the -35 region, partially overlapping the I_2_ half-site, was not inverted. For the fusion of sfGFP to the C-terminal of BLADE FP6, pBLADE(FP6*)-mCherry was amplified with primers 59 and 60, the *sfgfp* gene was amplified from pBLADE(FP6*)-sfGFP with primers 61 and 62, that carried a GS linker. The cloning yielded to pBLADE((FP6-sfGFP)*)-mCherry. Gibson Assembly was performed using NEBuilder® HiFi DNA Assembly (New England Biolabs). PCR were performed using the Phusion Flash High Fidelity PCR Master Mix (Thermo Scientific). Oligonucleotides were ordered at Sigma Aldrich.

To clone the extended library of BLADE TFs, we used a modular Golden Gate cloning strategy using an optimized junction set for part assembly taken from^85^. The overhangs as well as the individual parts and the final plasmid sequences are shown in Supplementary Table 6 as well as Supplementary Dataset 1. To invert the transcriptional unit containing the *mCherry* gene under AraC-controlled promoters, we first assembled the transcriptional unit separately, and then PCR-amplified the resulting fragment to create an A junction inverted at the end, and an F junction inverted at the beginning of the transcriptional unit and further treated the resulting construct as a part. Individual parts were first cloned into a part vector using BbsI-HF. The final plasmids were assembled from individual parts with BsaI-HF for digestion of the parts, and BbsI-HF for digestion of the plasmid backbone, which contains a p15a and a chloramphenicol acetyl transferase. Plasmids were transformed using a one-step preparation protocol of competent *E. coli* cells for transformation of plasmids in testing strains^86^. The sequences of all cloned plasmids were confirmed by Sanger sequencing (Eurofins Genomics Europe Sequencing GmbH, Köln, Germany, and Microsynth AG, Balgach, Switzerland). A list of all vectors used and constructed in this study is shown in Supplementary Table 4 and Supplementary Dataset 1. Oligonucleotide sequences used for PCR amplification and Golden Gate part sequences are shown in Supplementary Tables 5 and 6. The cloning was performed using chemically competent *E. coli* TOP10 cells (Thermo Scientific).

### Bacterial growth

For experiments shown in Figures 1-5 and Supplementary Figures 2-9, cultures were handled under safe red light whenever containing light-sensitive constructs. The cultures were incubated overnight in TB or LB (Figure 2 and Figure 3 g, h) medium and grown at 37°C (with the exception of cultures used for the experiments in Figure 4d and Supplementary Figure 8 which were grown at 18°C) in an incubator shaking at 250 rpm, in black plastic tube (Argos Technologies LiteSafe® 15 ml) if containing light-sensitive samples, in transparent glass tubes otherwise (with the exception of cultures used for the experiments in Figure 4d and Supplementary Figure 8 which were shacked at 110 rpm). The following morning, the cultures were diluted to OD_600_ 0.1 and let grow until OD_600_ 0.4. Half of the culture was then transferred in transparent glass tubes and induced either with blue light or with arabinose for 4 hours (with the exception of cultures used for the experiments in Figure 4d and Supplementary Figure 8 which were diluted 1:30). For experiments shown in Figure 6 and Supplementary Figures 12, and 14-17, cultures were grown in an environmental shaker. The shaking incubator consisted of a Kuhner ES-X shaking module (Adolf Kühner AG, Basel, Switzerland) mounted inside an aluminum housing (Tecan, Maennedorf, Switzerland) and temperature-controlled using an “Icecube” (Life imaging services, Basel, Switzerland). Cultures were grown at 37°C with shaking at 300 rpm in black, clear-bottom 96-well plates (Cell Culture Microplates 96 Well µClear® CELLSTAR®, Greiner Bio-One GmbH, Product #: 655090), which were sealed with peelable foil (Sealing foil, clear peelable for PlateLoc, No. 16985-001, Agilent) to prevent liquid evaporation and guarantee sterility, as well as a plastic lid (Greiner Bio-One GmbH, Product #: 656171). Overnight cultures were inoculated in M9 medium and grown over night to an OD_600_ of about 4. These cultures were diluted 1:20,000 into fresh M9 medium containing the respective inducer concentrations, right before the start of the experiment. This high dilution ensures that the cells are still in logarithmic growth phase after 5h, at the end of the experiment^61^. 200 µl of inoculated culture were incubated per well in the 96-well plates. Cells were grown for 5h before transcription and translation was stopped with rifampicin and tetracycline^61^. The inhibition solution contained 500 μg ml^-1^ rifampicin and 50 μg ml^-1^ tetracycline in phosphate buffered saline (Sigma-Aldrich Chemie GmbH, Dulbecco’s phosphate buffered saline) and was filtered using a 0.2 μm syringe filter (Sartorius). 100 µl inhibition solution were aliquoted in 96-well U-bottom plates (Thermo Scientific Nunc), precooled on ice and samples were added in equal volumes (100 µl), resulting in a final inhibitor concentration of 250 μg ml^-1^ rifampicin (Sigma-Aldrich Chemie GmbH) and 25 μg ml^-1^ tetracycline (Sigma-Aldrich Chemie GmbH). After sample was added, the solution was incubated on ice for at least 30 min. Then mCherry maturation was carried out at 37 °C for 90 min. The samples were kept at 4°C until measurement through flow cytometry.

### Light illumination systems

To illuminate the glass tubes in the shaker, six high-power 460 nm LEDs type CREE XP-E D5-15 (LED-TECH.DE) were used (Supplementary Fig. 18). The LEDs were connected to a power supply (Manson HCS-3102) that allowed to tune the voltage, hence the light intensity. Unless specified, the light intensity reaching the cultures was 5 W/m^2^ as measured with a LI-COR LI-250A Light Meter. For the bacterial photography and the induction of the library of genes with unknown or poorly defined function, we used a custom-made light box with, among others, 6 blue (455 nm) LEDs (Supplementary Fig. 19). To avoid generation of a blurred image in the bacteriograph, all the LEDs except for the one in the center were obscured with colored tape. The average light intensity reaching the plate was 5W/m^2^ with 6 LEDs and 1.3 W/m^2^ with one LED.

The 96-LED array was designed using CircuitMaker 1.3.0 (www.circuitmaker.com). The LEDs (SK6812, Dongguang Opsco Optoelectronics Co., Dongguan City, China) were arranged on the PCB at a pitch of 9 mm in an 8 x 12 grid to be compatible with standard 96-well plates. All LEDs were daisy-chained using their DIN and DOUT ports. A 0.1nF capacitor was placed in parallel to the VDD port of each LED as proposed by the manufacturer. The 2-layer circuit was manufactured on a 1.6 mm thick FR-4 substrate, and the surface of the PCBs was coated with black solder mask to reduce reflection. The PCBs were ordered preassembled with the LEDs and 0.1 nF capacitors (www.pcbway.com, Shenzhen, China). Every 96-LED PCB had one signal-in and one signal-out SMA connector such that several 96-LED PCBs could be daisy-chained using SMA cables and controlled by a single microcontroller. Up to 4x 96- LED PCBs could be powered using a single Adafruit #658 5V 10A switching power supply (digikey.ch, Munich, Germany) using a custom-made PCB to distribute the power to several LED arrays. The LEDs were controlled through an Arduino Uno microcontroller (Arduino, Somerville, MA, USA) using the fastLED library (http://fastled.io/).

The 96-LED array was mounted inside the shaking incubator using custom 3D-printed holders. The holders were printed with an Ultimaker S5 using black Ultimaker CPE (Ultimaker, Utrecht, Netherlands) to reduce reflections. For better dissipation and distribution of the heat generated by the LEDs, a custom-made anodized aluminum plate (10 mm thick, with 96 holes of 4 mm diameter) was mounted on top of the 96- LED array. Another 3D-printed adapter was placed between the aluminum plate and the microtiter plate to ensure optical insulation of the wells. The 3D-printed parts and the metal plate were aligned and held in place by metal rods (4 mm diameter, 20 mm length).

### Flow Cytometry

For experiments shown in Figures 1,3,4 and Supplementary Figures 3, 4, 5b, 9b and 9c, fluorescence was measured using the LSR Fortessa flow cytometer (BD Biosciences). Samples were centrifuged at 4000*g* for 4 min to remove the glycerol-containing solution, then the pellets were resuspended in PBS. Data analysis was performed using the open source FCSalyzer software. The mCherry fluorescence was excited with a 561 nm laser (50 mW), and emission was detected using a 610/20-nm filter pass (PMT voltage set to 750 V). The GFP fluorescence was excited with 488 nm laser (100 mW), and emission was detected using a 530/30-nm filter pass (PMT voltage set to 405 V). A forward scatter height (FSC-H) threshold of 1,400 was used to gate for living cells and eliminate debris. 10^5^ events per sample were recorded for each experiment. The cell density of the samples was manually regulated by addition of PBS in order to have less than 2*10^4^ events/s recorded by the machine. To compensate any variable that can alter the measurement of the fluorescence by the flow cytometer, each experiment was normalized with the fluorescence value of the negative control grown the same day of the experiment. For experiments shown in Figure 6 and Supplementary Figures 11, 12, 14, 15-17, fluorescence was measured on a Cytoflex S flow cytometer (Beckman Coulter) equipped with CytExpert 2.1.092 software. The mCherry fluorescence was excited with a 561 nm laser and emission was detected using a 610/20 nm band pass filter and following gain settings: forward scatter 100, side scatter 100, mCherry gain 3,000 when mCherry was expressed from the I_1_-I_2_ promoter, and 300 gain when mCherry was expressed from the I_1_-I_1_ promoter due to the difference in expression levels. Thresholds of 2,500 FSC-H and 1,000 SSC-H were used for all samples. The flow cytometer was calibrated before each experiment with QC beads (CytoFLEX Daily QC Fluorospheres, Beckman Coulter) to ensure comparable fluorescence values across experiments from different days. At least 15,000 events or 2 min were recorded in a two-dimensional forward and side scatter gate, which was drawn by eye and corresponded to the experimentally determined size of the testing strain at logarithmic growth and was kept constant for analysis of all experiments and used for calculations of the median and CV using the CytExpert software. The same gating strategy was previously used and is depicted in Supplementary Figure 21.

### Characterization of the FP1-FP7 VVD-AraC_DBD_ fusion constructs

Chemically competent *E. coli* MG1655 cells were transformed with pBLADE(FP1*/FP7*)- mCherry, pBLADE(FP1**/FP7**)-mCherry, pBLADE-empty (negative control), and pBLADE(AraC_WT_*/AraC_WT_**)-mCherry (positive controls). Overnight cultures of cells transformed with the FP1-FP7 fusions were diluted to OD_600_ 0.1, let grow in the dark to OD_600_ 0.4 and split into two cultures, one of which was kept in the dark and one of which was illuminated for 4 h. The overnight culture of the negative control was diluted to OD_600_ 0.1, and let grow for the same amount of time as all other cultures (circa 5 h 30 min). The overnight cultures of the positive controls were diluted to OD_600_ 0.1, let grow to OD_600_ 0.4 and split into two cultures, one of which was left without arabinose and one of which was induced with 0.1% arabinose for 4 h. After the induction time, 200 µl of each sample were collected, mixed with 200 µl of a transcription and translation inhibition solution (500 μg ml^-1^ rifampicin and 50 μg ml^-1^ doxycycline in phosphate buffered saline) and incubated in the dark 90 min at 37°C with 110 rpm shaking. This protocol allows obtaining a full maturation of almost all the mCherry proteins translated at the end of the induction time^1^. After the incubation with the inhibitor, samples were either diluted 10 times with PBS and immediately analyzed with the flow cytometer, or diluted 1:1 with 60% glycerol and frozen at −80°C.

### Dynamic control of gene expression

The overnight cultures transformed with pBLADE(FP6**)-mCherry and pBLADE-empty (negative control) were diluted in TB to OD_600_ 0.05 in dark tubes and let grow until OD_600_ 0.15. 200 µl of each sample were collected, mixed with 200 µl of a transcription and translation inhibition solution (500 μg ml^-1^ rifampicin and 50 μg ml^-1^ doxycycline in phosphate buffered saline), incubated in the dark 90 min at 37°C with 110 rpm shaking, diluted 1:1 with 60% glycerol, and frozen at −80°C. The rest of the culture was transferred in a transparent glass tube and illuminated with blue light as described (Light illumination systems) for 2 h. Then, another aliquot was taken and frozen, and the remaining culture was diluted to OD_600_ 0.15 again and transferred to a dark tube, for a total of three dark-light cycles.

### Measurement of the kinetics of BLADE-mediated mCherry expression

Chemically competent *E. coli* MG1655 cells were transformed with pBLADE(FP6*)- mCherry and pBLADE-empty. The overnight cultures were diluted and each split into two cultures, of which one was induced with blue light and one kept in the dark. Every hour for 6 h, 200 µl of each sample were collected, mixed with 200 µl of the transcription and translation inhibition solution, incubated in the dark 90 min at 37°C with 110 rpm shaking, diluted 1:1 with 60% glycerol, frozen at −80°C and subsequently analyzed with the flow cytometer.

### Light intensity titration

Chemically competent *E. coli* MG1655 cells were transformed with pBLADE(FP6**)-mCherry and pBLADE-empty. The overnight culture of the cells transformed with pBLADE(FP6**)-mCherry was diluted and split into 5 independent cultures, each of which was induced with blue light of different intensity (which was tuned adjusting the voltage in the power supply connected to the LEDs) for 4 h. The overnight culture of the cells transformed with pBLADE-empty was diluted and grown in the dark for 4 h. 200 µl of each sample were then collected, mixed with 200 µl of the transcription and translation inhibition solution, incubated in the dark 90 min at 37°C with 110 rpm shaking, diluted 1:1 with 60% glycerol, frozen at −80°C and subsequently analyzed with the flow cytometer.

### Bacterial photography

Chemically competent *E. coli* MG1655 cells were transformed with pBLADE(FP6*)-sfGFP. The overnight culture was diluted in LB to OD_600_ 0.1 and grown for approximately 6 h. A 96-well lid (12.7 x 8.5cm) was filled with 30-40 ml of 1% LB-agar and let solidify. 1 ml of the culture was then mixed with 9 ml of 0.4% agar at 42°C (measured with infrared thermometer TFA Dostmann (Wertheim-Reicholzheim, Germany)) and plated on top of the solidified agar in the 96- well lid. The plate was covered with a transparent plexiglass parallelepiped with the Blade Runner movie poster sticker. To increase the opacity of the dark zones of the picture, three identical stickers were overlapped on one another. The plate was then placed in a 37°C incubator under the light box overnight. The next morning the plate was imaged with a Zeiss Axio Zoom.V16 stereo zoom microscope equipped with PlanNeoFluar Z 1.0x objective, zoom 0.7x, AxioCam MR R3 camera and the 38 HE filter set (Ex BP 470/40, FT 495, Em BP 525/50; sfGFP). The bacteriograph is composed of 110 tiles stitched together with ZEN Blue software.

### DIC and fluorescence microscopy

5 µl of the bacterial culture were applied to a thin agarose pad composed of 1% agarose for microscopy at room temperature and of 1% agarose and 0.1% LB in Tethering buffer (10 mM potassium phosphate, 0.1 mM EDTA, 1 mM L-methionine and 10 mM sodium lactate; pH 7.0) for long-term microscopy at 37°C. Images were acquired on a Zeiss Axio Observer Z1/7 fluorescence microscope equipped with an Alpha Plan-Apochromat 100x/1.46 Oil DIC (UV) M27 objective, filter sets 38 HE (Ex BP 470/40, FT 495, Em BP 525/50; sfGFP), 108 HE (Ex BP 423/44, DBS 450+538, Em DBP 467/24+598/110; MM 4-64), 96 HE (Ex BP 390/40, FT 420, Em BP 450/40; DAPI), 64 HE (Ex BP 587/25, FT 605, Em BP 647/70; mCherry) and an Axiocam 506 Mono camera. To image the library of genes with unknown or poorly defined function in a fast and efficient way, the samples (circa 5 ml) were applied to a 96-well lid, which was filled with 1% agarose, let solidify and covered with two 75 x 50 mm glass coverslips (Carl Roth GmbH, Karlsruhe). Before imaging, samples were incubated for 5 min with 1.2 µg ml^-1^ of the membrane dye MM 4-64 (AAT Bioquest Sunnyvale, CA) and 0.5 µg ml^-1^ of 4′,6-diamidino-2-phenylindole (DAPI, Sigma-Aldrich Chemie GmbH).

The induction of gene expression in selected cells within a population of MG1655 cells transformed with pBLADE(FP6**)-sfGFP was performed on a Zeiss LSM 800 confocal microscope. An area of 6.4 µm^2^ was illuminated with a 488 nm diode laser (10 mW) at 0.1% intensity, with a frame average of 8, resulting in 0.36 µs of light per pixel. The illumination was given in pulses of 5 min for a duration of 3 h.

### FRAP

FRAP was performed on a Zeiss LSM 800 confocal microscope. An overnight culture of MG1655 cells transformed with pBLADE((FP6-sfGFP)*)-mCherry was diluted in the morning in fresh TB medium to OD_600_ 0.1, and grown until it reached OD_600_ 0.4. 5 µl of the culture were then applied to a thin 1% agarose pad. After selecting a cell with a bright fluorescent spot, the area to bleach within the cell (whole spot) was manually set (the whole focus) and bleached with a single 1 s pulse of a 488 nm diode laser (10 mw) at 50% intensity. An image in the GFP channel (filter set 38 HE: Ex BP 470/40, FT 495, Em BP 525/50) was taken 5- and 15-min post bleaching to measure the recovery of the fluorescent signal.

### Induction of *rodZ* in KC717 cells

Strain KC717 (kind gift of KC Huang, Stanford University) was grown in LB medium supplemented with 0.2% arabinose (to maintain the cells rod-shaped) during transformation of chemically competent KC717 cells and DNA extraction procedures. The blue light and arabinose induction were performed as described above. The recovery phase of the culture induced with arabinose was performed by centrifuging it at 6000*g* for 4 min and resuspending it with the same volume of LB. The centrifugation and resuspension steps were repeated a second time and the culture was then diluted to OD_600_ 0.1. The recovery phase of the culture transformed with pBLADE(FP4**) was performed by dilution of the culture exposed to blue light OD_600_ 0.1 and incubation in the dark.

### BLADE (FP6) expression and purification

Chemically competent *E. coli* Rosetta (DE3) cells carrying the pLysS plasmid were freshly transformed with pET28a-FP6 and cultivated overnight in LB medium supplemented with 50 μg ml^−1^ kanamycin. LB medium (1 l) containing kanamycin was inoculated using the pre-culture to obtain OD_600_ of 0.1. The culture was grown at 37°C until OD_600_ of 0.5, after which 1 mM IPTG and 5 μM FAD were added, and the culture was grown for 16 h at 18°C under constant blue light. Cells were collected by centrifugation and the pellet was re-suspended in 30 ml of lysis buffer (50 mM potassium phosphate pH 8.0, 300 mM NaCl and 10 mM imidazole pH 8.0) supplemented with a cOmplete™ protease inhibitor cocktail tablet (Roche). Cell lysis was performed by sonication and the lysate was centrifuged at 20,000 rpm for 20 min at 4°C. The supernatant was then co-incubated with 1 ml of HisPur™ Ni-NTA Resin (Thermo Scientific) for 2 h at 4°C. Protein purification was performed by the gravity flow method. The bound proteins were washed twice with 5 ml of wash buffer (lysis buffer + 10 % glycerol + 20 mM imidazole) and finally eluted with 1.5 ml of elution buffer (50 mM potassium phosphate pH 7.5, 300 mM NaCl, 500 mM imidazole pH 8.0 and 10 % glycerol). The elution buffer was replaced with a storage buffer (20 mM HEPES-NaOH pH 7.5, 150 mM NaCl and 10 % glycerol) using an Amicon® Ultra-4 regenerated cellulose NMWL 10 kDa centrifugal filter unit (Merck). The protein was then stored as 50 μl aliquots at −80°C. We verified that the purified protein could respond to light by measuring the absorption spectrum (Supplementary Fig. 20).

### Spectroscopy

The absorption spectrum of the FAD cofactor bound to VVD within BLADE (FP6) was measured exciting the sample in the 300-600 nm range using a Multiskan GO (Thermo Scientific) plate reader. The protein sample was incubated 4 days at 4°C in the dark in a buffer solution (25 mM HEPES, 150 mM NaCl, 10% glycerol, 0.1% EDTA; pH 7.5) and then diluted to 0.5 mg ml^-1^ prior to the measurement of the absorption spectrum in the dark state. The same sample was then illuminated with blue light (455 nm; 50 W/m^2^) for 5 min at room temperature and the absorption spectrum in the lit state was measured. The absorption spectrum of the blank (only medium) was subtracted from the dark and lit state spectra.

### SEC

Purified BLADE (FP6) was thawed and stored in complete darkness at 4°C for 6 days. The sample (1 ml of protein with a concentration of 0.5 mg ml^-1^) was loaded onto a Superdex™ 75 Increase 10/300 GL (GE Healthcare Lifesciences) column at 4°C. The running buffer consisted of 20 mM HEPES-NaOH pH 7.5, 150 mM NaCl and 10 % glycerol, and the flowrate was adjusted to 0.25 ml min^-1^. Dimerization of BLADE FP6 was triggered by incubating the protein under constant blue light (455 nm; 50 W/m^2^) for 30 min at 4°C, prior to injection. During the run, the column was either illuminated with constant blue light (460 nm; 8 W/m^2^, lit sample) or kept in complete darkness (dark sample). Bovine serum albumin (BSA) and carbonic anhydrase (CA) were used as size markers at a concentration of 0.5 mg ml^-1^ each.

### Light-induced expression of genes with unknown or poorly defined function

Chemically competent MG1655 cells were transformed with the 117 pBLADE plasmids constituting the library to characterize the 39 genes with unknown or poorly defined function. Cultures were grown in the dark overnight in LB in non-treated 96- well plate (VWR, Radnor, PA) at 37°C with 110 rpm shaking. The following morning a Scienceware® replicator (96-well; Merck KGaA, Darmstadt, Germany) was used to transfer about 5 μl of each culture into a fresh 96-well plate with 145 μl of TB in each well. The diluted cultures were incubated at 18°C with 110 rpm shaking for 1 h in the dark and then they were induced with blue light (455 nm, 5 W/m^2^) for 4 h.

### Measurement of bacterial growth

The growth curves of the cells transformed with the library of genes with unknown or poorly defined function were measured on a Synergy H4 Hybrid plate reader (BioTek) in 96-well plates. The cultures were grown in the dark overnight in LB in a 96-well plate at 37°C with 110 rpm shaking. The following morning the cultures were diluted to OD_600_ 0.1 in a fresh 96-well plate with 120 μl of LB. To prevent evaporation of the medium, also the unused wells of the plate were filled with the same amount of LB and the lid was sealed with parafilm. The plate was then illuminated with blue light (460 nm) and the OD_600_ of the culture was measured every 2 min in constant shaking for 20 h.

The overnight cultures of three selected members of the library (y*daT, ydhL, yhcF)* were diluted in LB to OD_600_ 0.1 and grown until they reached OD_600_ 0.4. Each culture was then split into two tubes, one of which was kept in the dark and one of which was illuminated for 4 h at 37°C with shaking at 250 rpm. The OD_600_ was measured at the end of the growth with the OD600 DiluPhotometer™ (Implen).

### Quantification of cell length, width and roundness

The cell length and width were calculated by first staining the cell with the membrane dye MM 4-64 (AAT Bioquest Sunnyvale, CA) to visualize the cell contour, and then manually measuring the long and short axes of the cell, respectively, using the straight-line ‘Selection’ tool of Fiji. At least 500 cells were measured for each sample. The histograms were generated in Excel by the Analysis ToolPak’s Histogram option. The roundness was manually calculated with the oval ‘Selection’ tool on unstained cells, using Fiji. At least 200 cells were measured for each sample.

### Computational prediction of function and localization of 39 genes with unknown or poorly defined function

We randomly selected 34 genes out of the y-ome, defined as the group of genes lacking to date experimental evidence of function^36^. We manually checked that the selected genes were not mentioned in any publication using several search engines. As controls for our pipeline, we included 5 genes for which some information was available (*ydaT*^37^; *ydiY*^38^; *ycbK* (MepK)^39^; *yehS*^40^; and *yebE*^41, 42^). We retrieved the amino acid sequences of the proteins encoded by all 39 genes in FASTA format and submitted them to the following webservers: Argot2.5^46^, PANNZER2^47^, DeepGoPlus^48^ and Phyre2^49^. The consensus localization and function were calculated as the output provided by at least 2/4 prediction tools.

### Mathematical modelling

The LacI IPTG dose-response was fitted to a Hill equation of the following form:

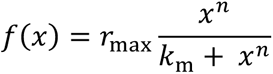

where *f(x)* describes the gene expression controlled by LacI, *x* represents the IPTG concentration, *r*_ma*x*_ is the maximal promoter expression, *k_m_* is IPTG’s dissociation constant for LacI, and *n* is the Hill coefficient for LacI. This dose-response was used subsequently to obtain IPTG concentration estimates from the fluorescence readouts of the constitutive promoters. All data were fitted using a non-linear least squares optimizer (MATLAB, MathWorks) with fitted parameter values r_max_ = 21352, k_m_ = 62, n = 1.7.

## Acknowledgements

We thank Maximilian Hörner for his help with the determination of the absorption spectrum of BLADE, João Nuno de Sousa Machado for his help with size-exclusion chromatography, Yanik Weber for help with characterizing the 96-well light induction plate, KC Huang for sharing with us KC717 strain, and Stephanie Aoki for helpful discussions. This study was funded by the DFG (grant no. VE776/2-1 to B.D.V.), by the BMBF (grant no. 031L0079 to B.D.V.), by the Excellence Initiative of the German Federal and State Governments BIOSS (Centre for Biological Signalling Studies; EXC-294), by the European Research Council (ERC-Advanced) under the European Union’s Horizon 2020 research and innovation programme (grant agreement number 743269).

## Author contributions

B.D.V. and A.B. conceived the study. B.D.V and M.K. supervised the study, and secured funding. E.R., A.B., E.A., N.P., M.K. and B.D.V. designed experiments and interpreted the data. E.R., A.B., M.H. and E.A. performed *in vivo* experiments. N.P. purified BLADE, and performed size-exclusion chromatography. L.E. performed initial experiments, which validated the idea. G.S. developed the 96-well light setup in collaboration with A.B. M.A.Ö. performed bioinformatics and computational structural biology analyses of the genes with unknown function. E.R., A.B., M.K. and B.D.V. wrote the manuscript.

## Data availability

The plasmids constructed in this study will be deposited on Addgene and will be additionally available from the corresponding authors upon request. The raw data supporting the conclusions of the paper will also be available from the corresponding authors upon request.

