## Supplementary Information for "An inducible AraC that responds to blue light instead of arabinose"

Romano, Baumschlager *et al.*

Table of contents:


- Supplementary Figures 1-21
- References

```
             -35                               -10
J23101    5’-TTTACAGCTAGCTCAGTCCTAGGTATAATGCTAGC - 3’
J23101*   5’-TTTACA-CTAGCTCAGTCCTAGGTATA-TGCTAGC - 3’
J23101**  5’-TTTACAGCTAGCTCAGTCCTAGGTATA-TGCTAGC - 3’
```

**Supplementary Fig. 1| Constitutive promoters used to drive the expression of the chimeric VVD-AraC fusion proteins.** Alignment of the nucleotide sequences of the original J23101 promoter[1] and the two variants that spontaneously arose in *E. coli* TOP10 cells after transformation of the pBLADE(FP4)-mCherry construct. The -35 and -10 regions are indicated with a white box for JS23101.

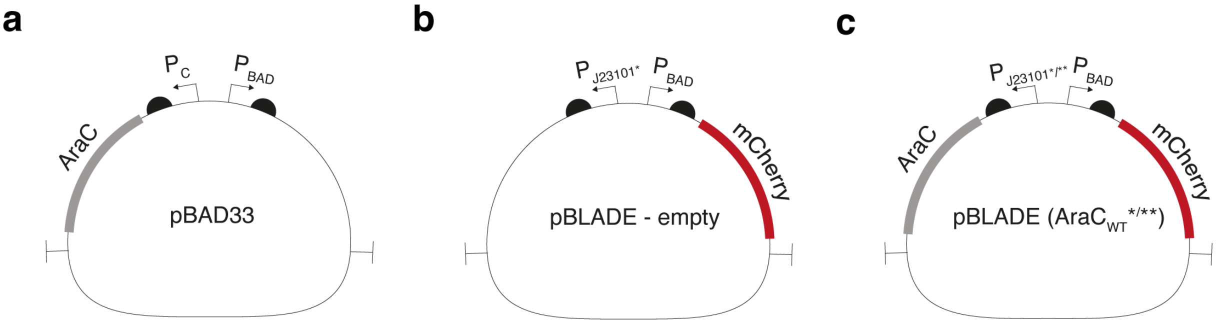


**Supplementary Fig. 2| Simplified maps of the original pBAD33 and the negative and positive control derivatives used in this study.** In pBAD33, the transcription factor AraC is transcribed from the wild type $P_C$ promoter, which is subject to negative feedback from AraC. The negative control pBLADE-empty contains the $P_{BAD}$ promoter driving the expression of mCherry but no transcription factor. The positive controls pBLADE(AraC$_{WT}$*)-mCherry and pBLADE(AraC$_{WT}$**)-mCherry are nothing but pBAD33 with the constitutive promoters J23101* and J23101** in place of $P_C$ to drive the expression of AraC.

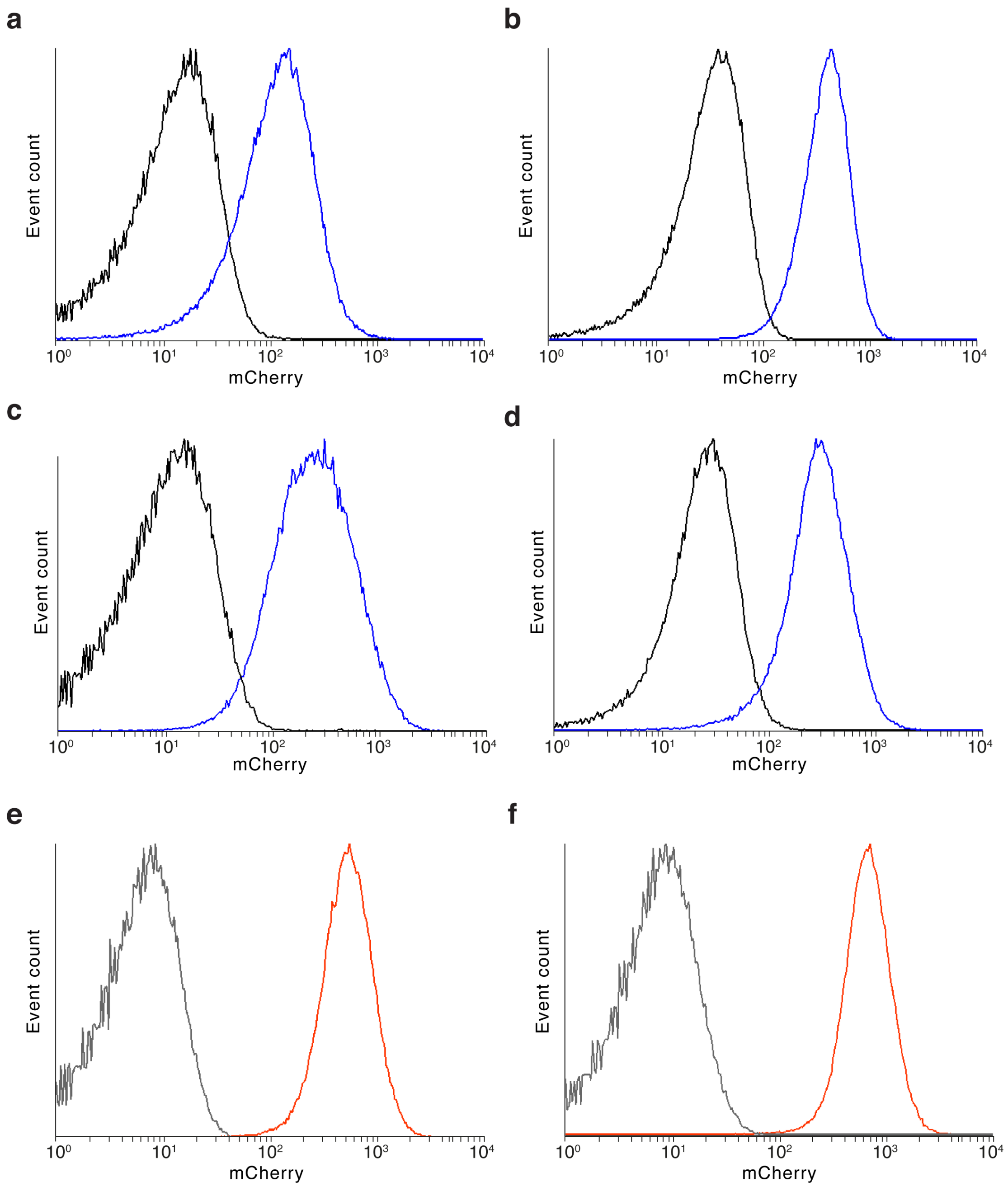


**Supplementary Fig. 3| Examples of histograms showing the distribution of the mCherry fluorescence within a population of bacterial cells. a-d**, MG1655 cells transformed with pBLADE(FP4*)-mCherry (**a**), pBLADE(FP4**)-mCherry (**b**), pBLADE(FP6*)-mCherry (**c**) and pBLADE(FP6**)-mCherry (**d**) were analyzed by flow cytometry after 4 h of blue light illumination (460 nm, 5W/m$^2$; blue curve) or 4 h in the dark (black curve). **e**,**f**, MG1655 cells transformed with pBLADE(AraC$_{WT}$*)-mCherry (**e**) and pBLADE(AraC$_{WT}$**)-mCherry (**f**) were analyzed by flow cytometry after 4 h of induction with 0.1% arabinose (red curve) or 4 h in the absence of arabinose (black curve).

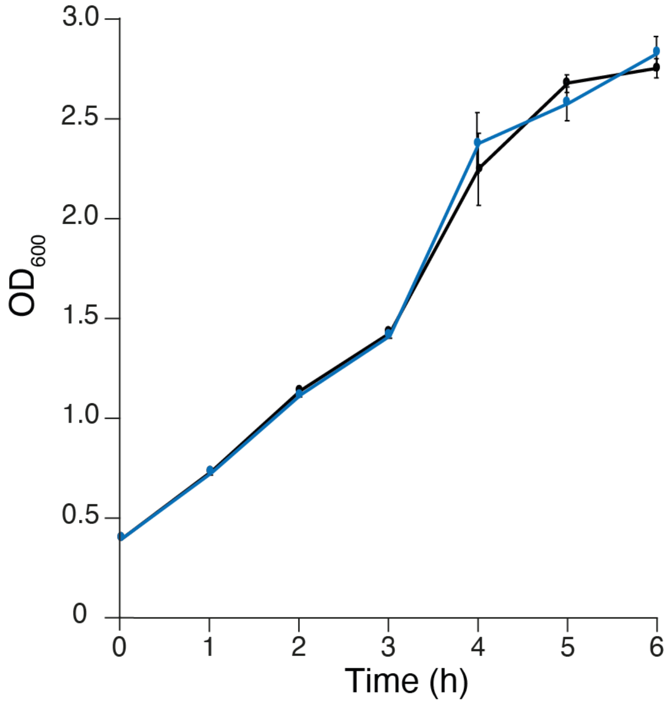


**Supplementary Fig. 4| Blue light is not toxic to the bacterial cells.** Growth curves of *E. coli* MG1655 cells transformed with pBLADE-empty illuminated with blue light (5 W/m$^2$; blue curve) or kept in the dark (black curve). Samples were collected every 60 min.

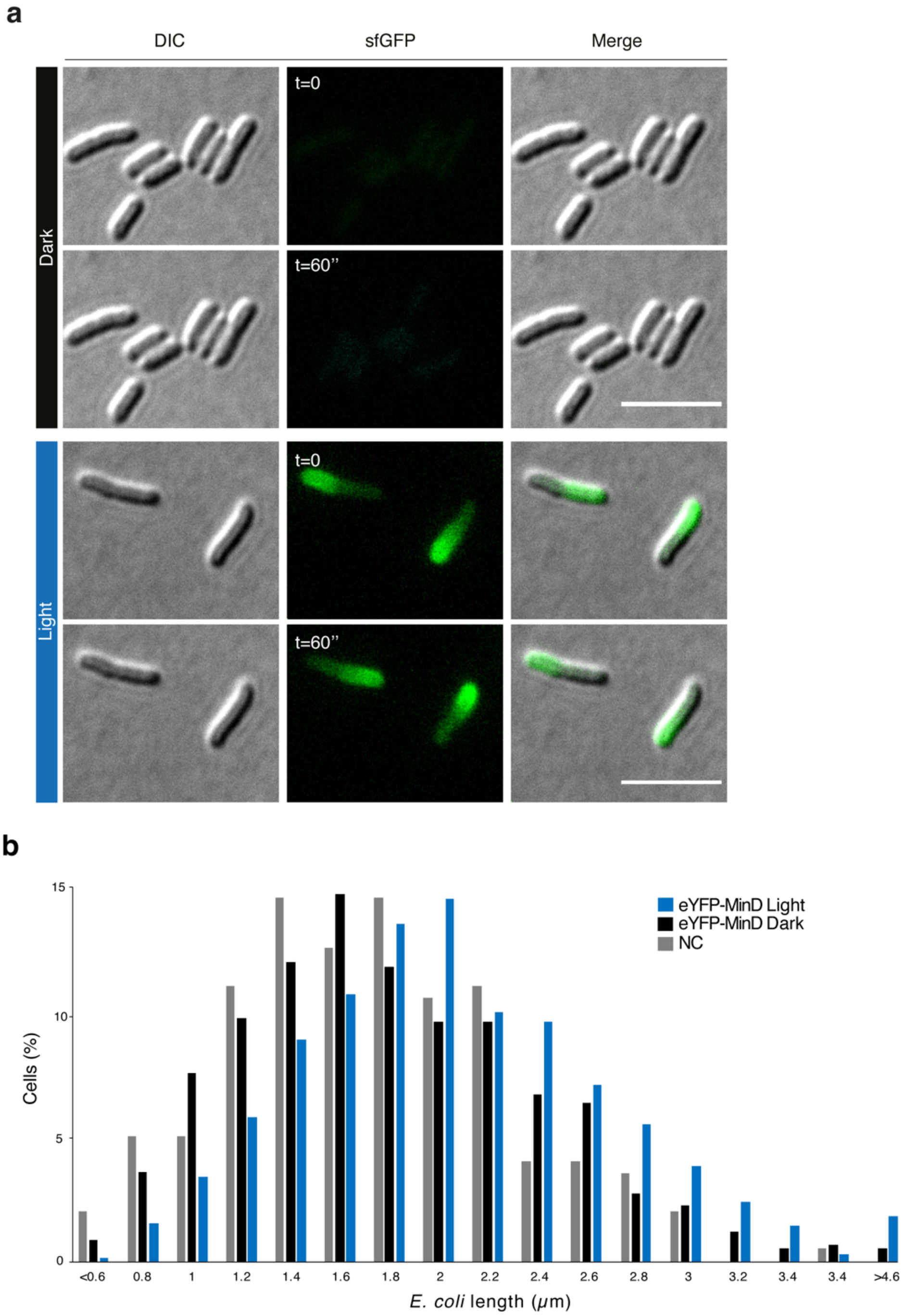


**Supplementary Fig. 5| BLADE can be used to tightly control the expression of functional bacterial proteins.** **a**, *E. coli* MG1655 cells transformed with pBLADE(FP6*)-eYFP-MinD were illuminated with blue light (5 W/m$^2$) for 2 hours or kept in the dark for the same time and imaged at the fluorescence microscope. The images show two snapshots of a time-lapse microscopy experiments at the indicated time points. Scale bar, 5 μm. **b**, Quantification of the cell length distribution for MG1655 cells transformed with the indicted constructs treated as in (**a**). The negative control (NC) was left in the dark.

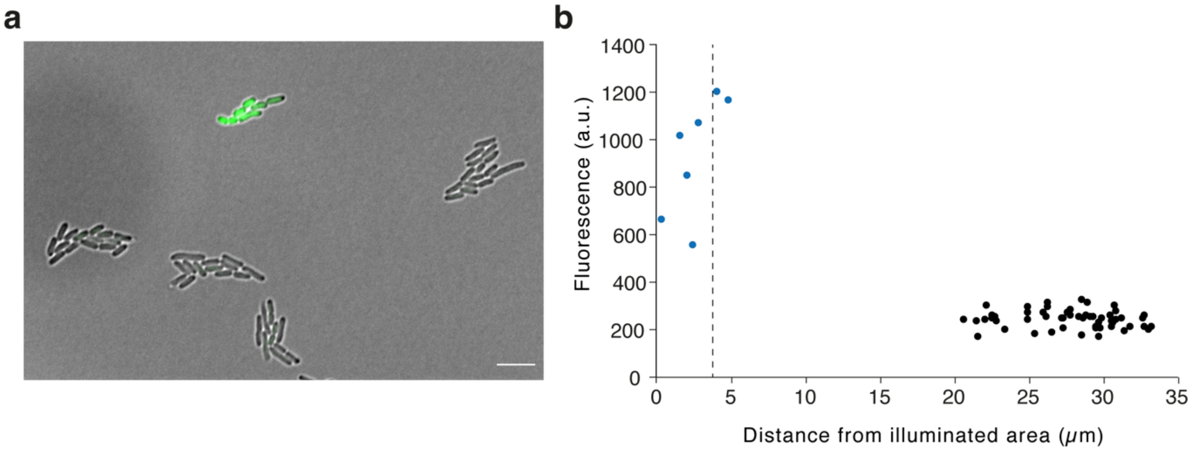


**Supplementary Fig. 6| Spatial control of gene expression. a**, Representative microscopy image showing the overly of the DIC and GFP fluorescence channel of *E. coli* MG1655 cells transformed with pBLADE(FP6**)-sfGFP taken after 3 h of blue light illumination of the area indicated by the blue box. Scale bar, 5 μm. **b**, Quantification of the GFP fluorescence of single cells within the illuminated microcolony (blue circles) and within non-illuminated microcolonies (dark circles) as a function of their distance from the center of the illuminated area (dashed line).

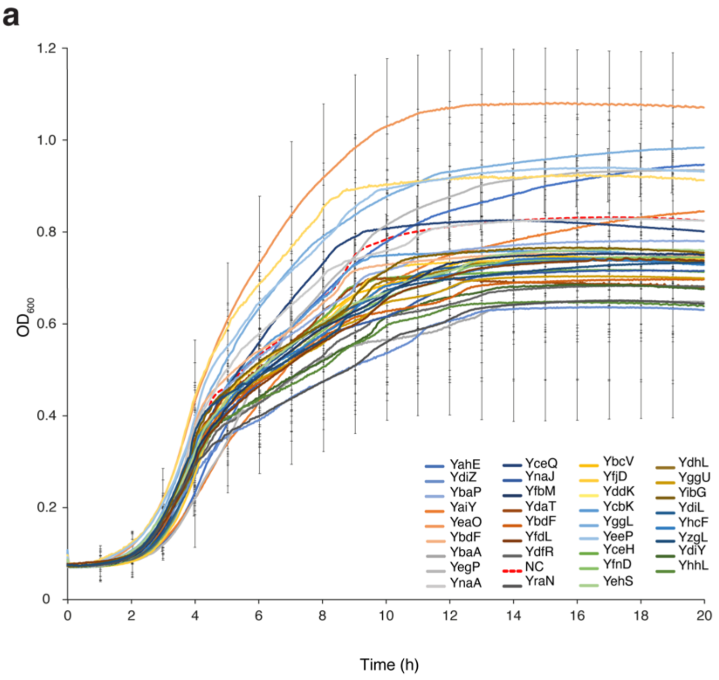


**b**

| Protein Name | $OD_{600}$ dark | $OD_{600}$ light |
|---|---|---|
| YdaT | 3.3 | 3.2 |
| YdhL | 3.2 | 3.0 |
| YhcF | 3.0 | 1.3 |
| NC | 3.8 | 3.8 |

**Supplementary Fig. 7| Most analyzed genes with unknown or poorly defined function do not alter the growth of bacterial cells. a**, Growth curves of *E. coli* MG1655 cells transformed with pBLADE carrying the genes coding for the indicated proteins for which the two-tailed, homoscedastic Student's *t* test resulted in non-significant growth alteration from the cells transformed with the empty plasmid (NC). Values represent mean ± s.d. of $n=3$ independent experiments. **b**, $OD_{600}$ of the indicated proteins grown in individual tubes in an incubator for 3 hours in the dark or under 460 nm light (5 $W/m^2$) illumination. Values represent a single experiment.

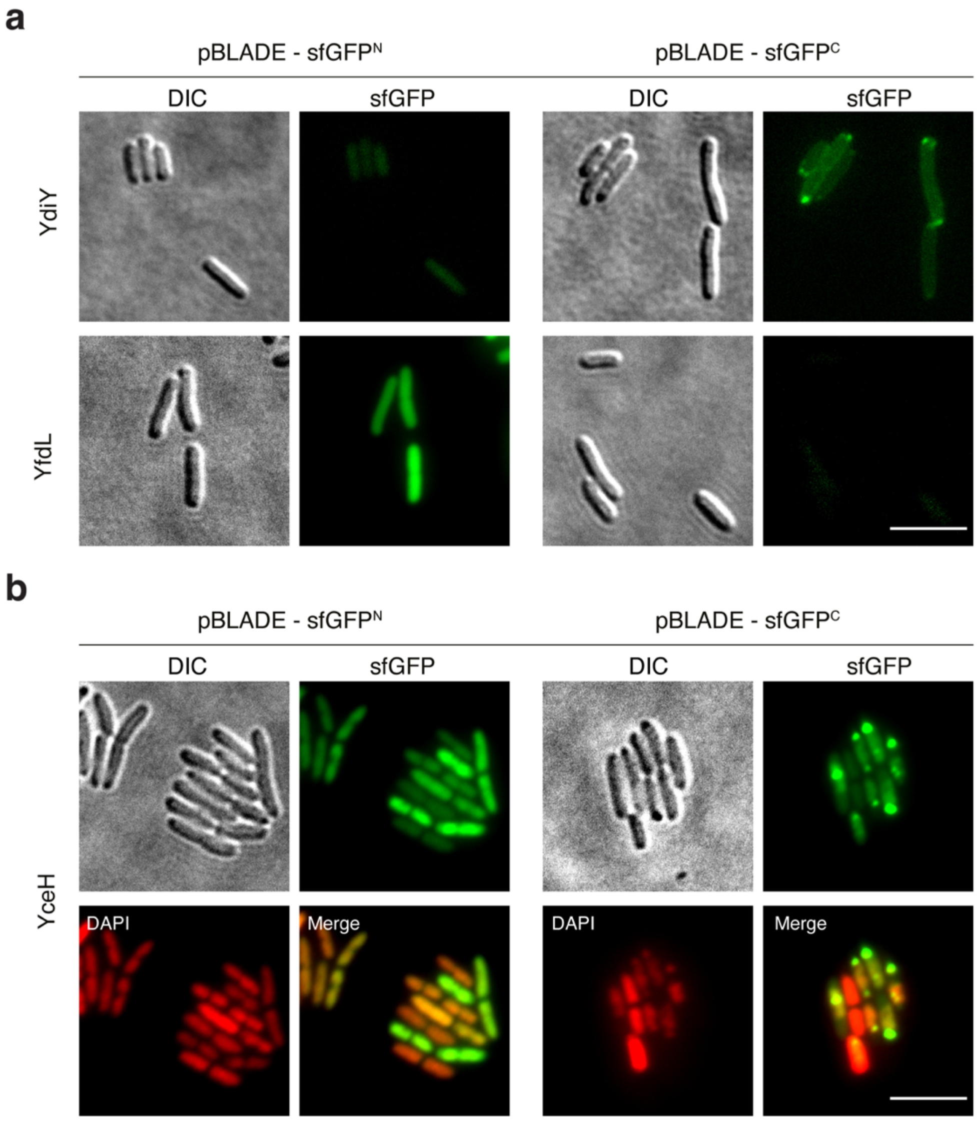


**Supplementary Fig. 8| The presence of the fluorescent protein sfGFP may alter the expression and localization of a gene of interest. a**,**b**, Representative images of *E. coli* MG1655 cells transformed with the indicated constructs grown for 4 h under 455 nm light illumination (5 W/m$^2$). pBLADE-sfGFP$^N$, N-terminal fusion; pBLADE-sfGFP$^C$, C-terminal fusion. Scale bar, 5 μm.

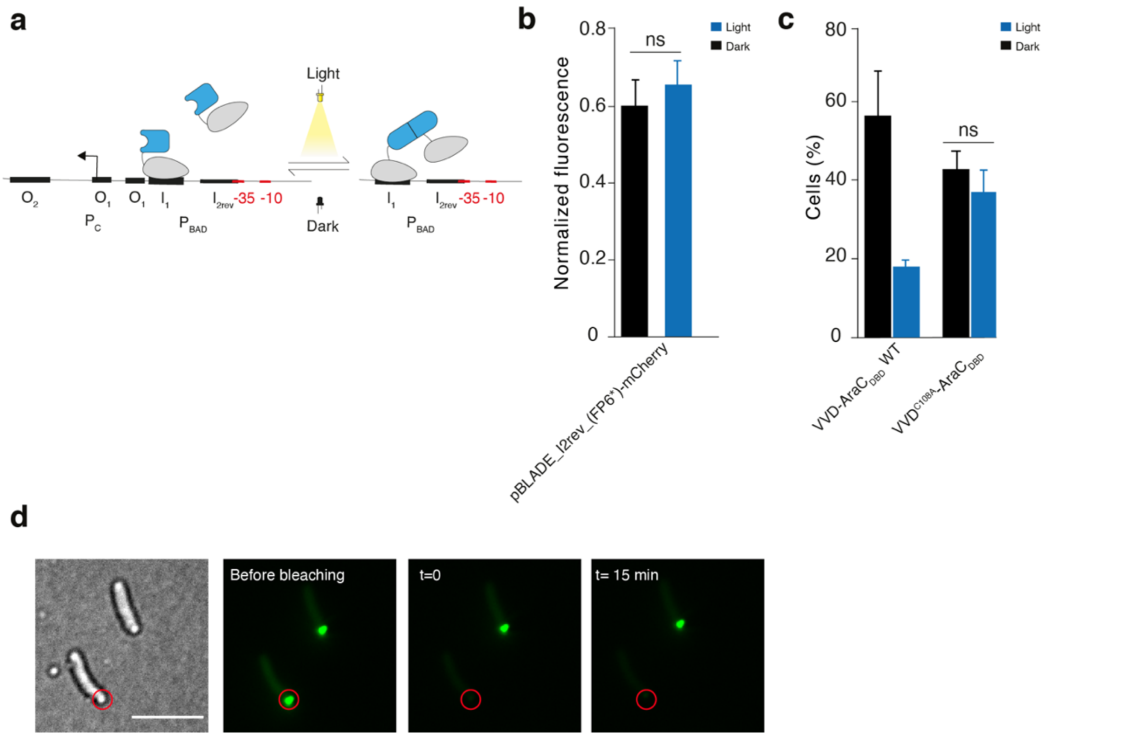


**Supplementary Fig. 9| The mechanism of BLADE-mediated gene expression involves the binding of the $I_2$ half-site in the light and the formation of aggregates in the dark. a**, Schematic representation of the synthetic promoter with the inverted $I_2$ half-site. The expected outcome is that dimeric BLADE fails to bind to this site and, therefore, to recruit the RNA polymerase. **b**, MG1655 cells transformed with pBLADE_$I_{2rev}$_(FP6*)-mCherry grown 4 h either in the dark or under 460 nm light (5 $W/m^2$) illumination were analyzed by flow cytometry. The fluorescence values were normalized to those of the negative control (cells transformed with pBLADE-empty). **c**, Quantification of the number of cells showing aggregates for the indicated constructs and conditions. **d**, Fluorescence does not recover after photobleaching of a BLADE-sfGFP aggregate. Scale bar, 5 µm.

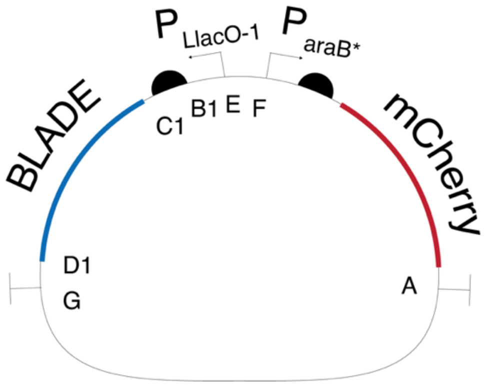


**Supplementary Fig. 10| Modular single plasmid.** Simplified map of the plasmid used to express the chimeric BLADE transcription factors (cTFs; for simplicity here indicated as BLADE) under the IPTG-inducible PLlacO-1 promoter. $P_{araB*}$, $P_{BAD}$ promoter deprived of all upstream regulatory sequences ($O_1$ and $O_2$ half-sites). Inside the plasmid overhang sites used for cloning as indicated in Supplementary Table 6 are shown.

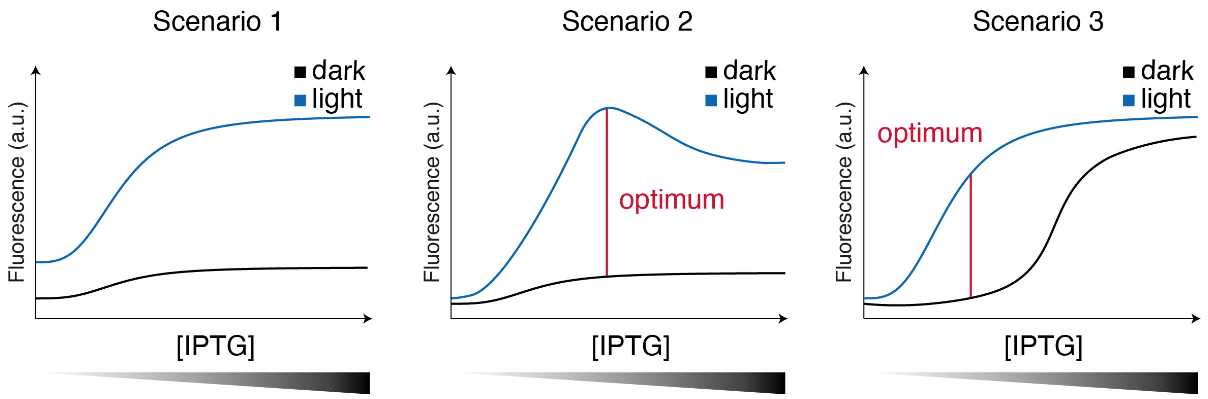


**Supplementary Fig. 11| Optimal gene expression output may be obtained at intermediate TF concentrations.** Schematic representation of different scenarios of dark and light-induced gene expression at different concentrations of the optogenetic TF, which is controlled via IPTG induction.

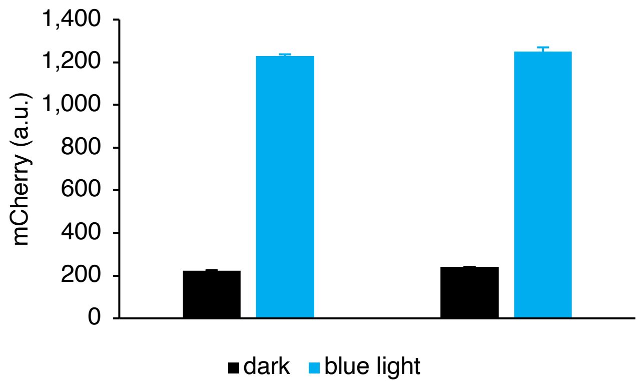


**Supplementary Fig. 12| Endogenous AraC does not influce the performance of BLADE TFs.** Comparison of functionality of VVD::$G_6$S::$\mathrm{AraC_{DBD}}$ in SKA684, the testing strain used in this study in which *araC* was deleted from the genome, and in AB360[1], which still contains a genomic copy of wild-type *araC*. Values represent mean ± s.d. of three independent experiments.

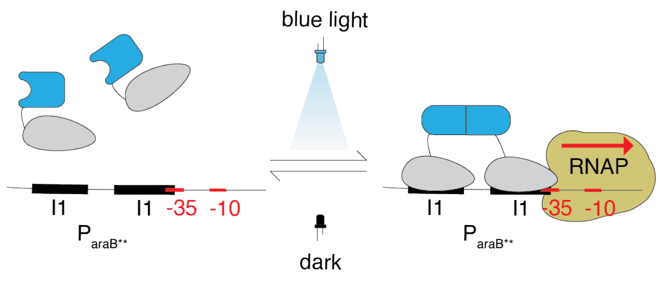


**Supplementary Fig. 13| Synthetic $\mathrm{P_{BAD}}$ promoter used in this study.** Schematic representation of the synthetic promoter containing two copies of the $I_1$ half-site and the hypothesized mode of action.

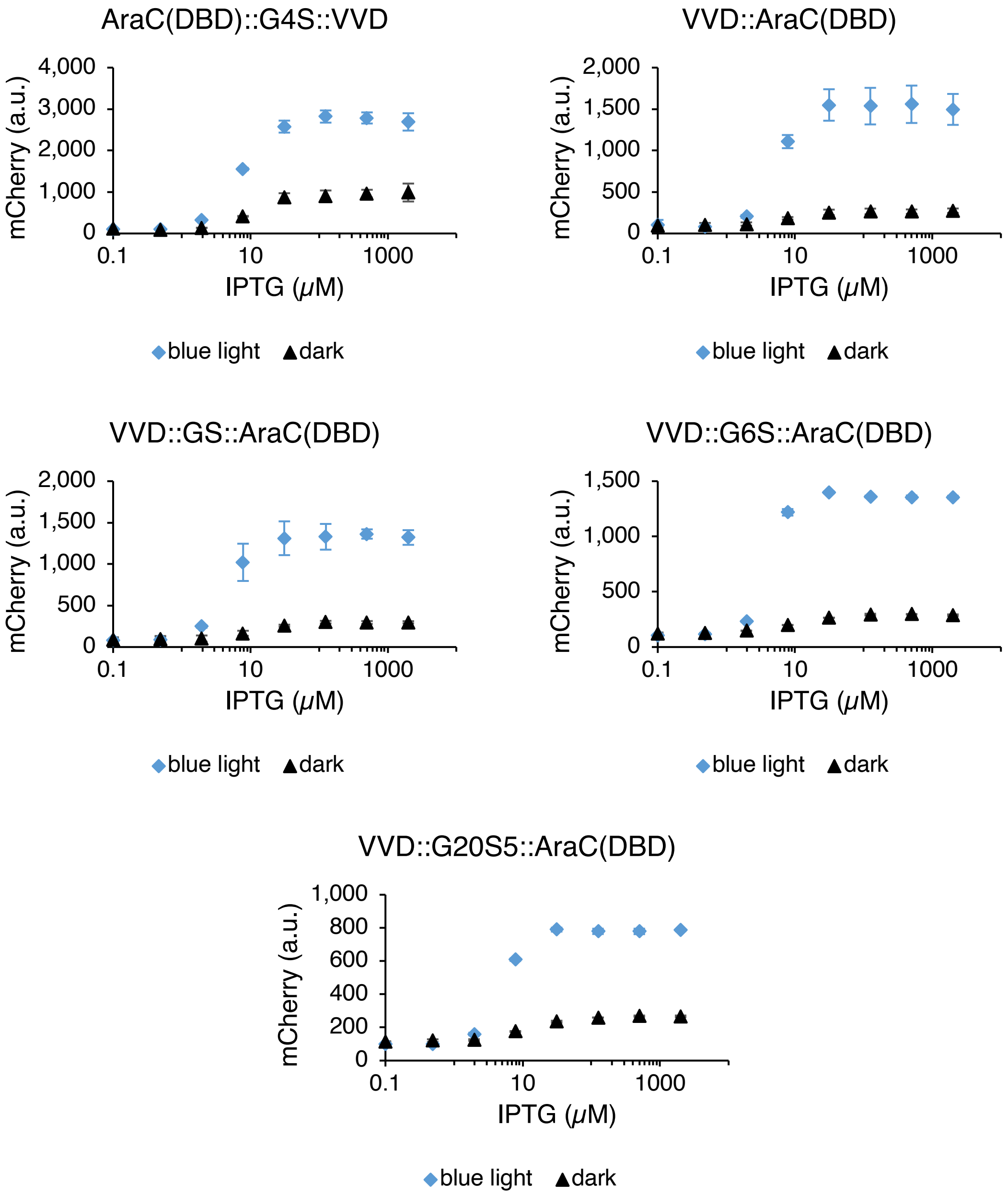


**Supplementary Fig. 14| Examples of IPTG-dose response curves for the VVD-AraC chimeric constructs.** IPTG-dose response curves of cells transformed with the single plasmid bearing the indicated VVD-AraC fusions and mCherry under control of $P_{BAD}$ deprived of all upstream regulatory elements. Cells were either in the dark, or induced with saturating blue light. Values represent mean ± s.d. of either triplicates or duplicates for mCherry fluorescence.

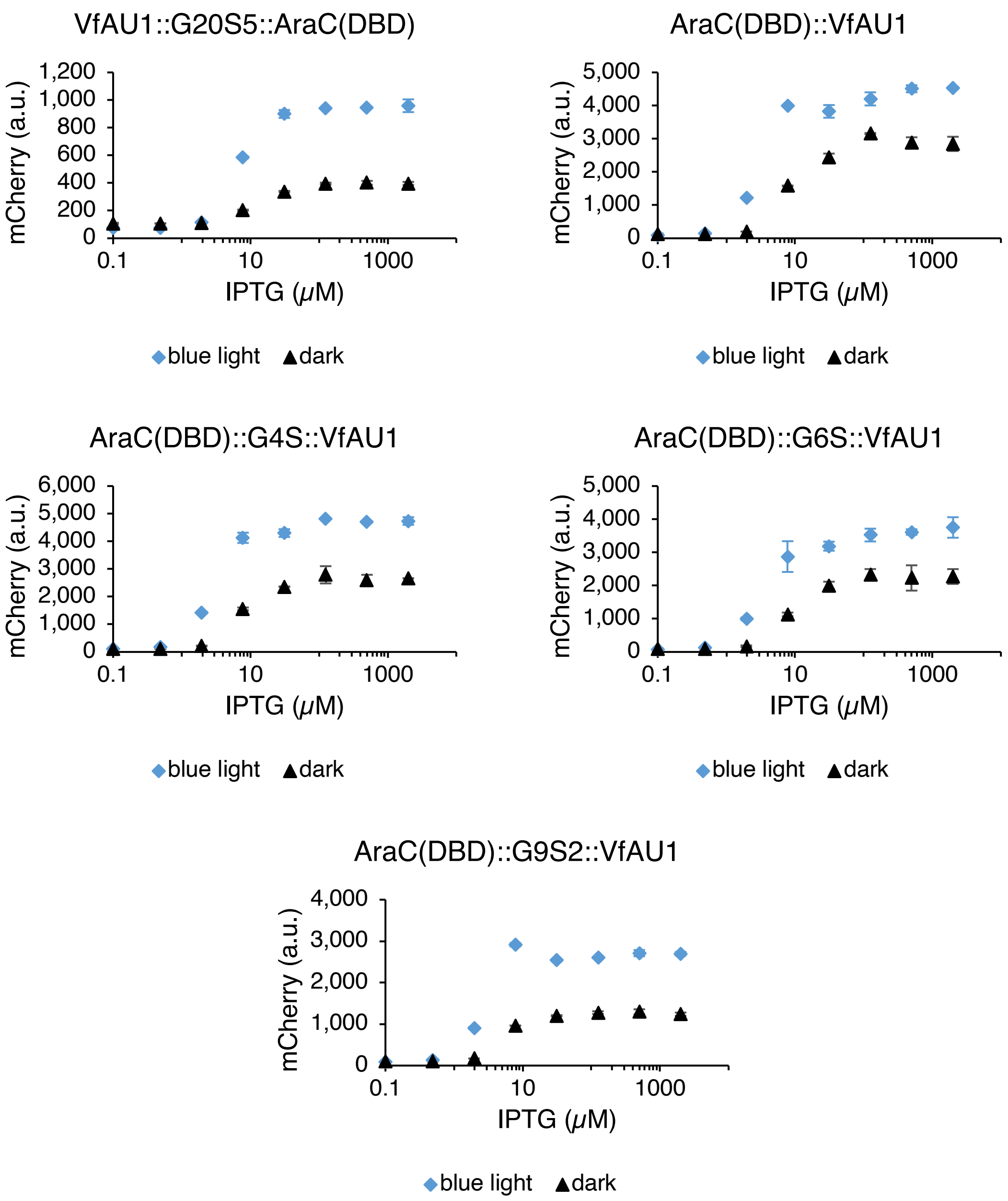


**Supplementary Fig. 15| Examples of IPTG-dose response curves for the VfAu1-AraC chimeric constructs.** IPTG-dose response curves of cells transformed with the single plasmid bearing the indicated VfAu1-AraC fusions and mCherry under control of $P_{BAD}$ deprived of all upstream regulatory elements. AraC(DBD)::G20S5::VfAu1 contained the RBS RiboJ+B0033m, while VfAu1::G20S5::AraC(DBD) contained the RBS 116. Cells were either in the dark, or induced with saturating blue light. Values represent mean ± s.d. of either triplicates or duplicates for mCherry fluorescence.

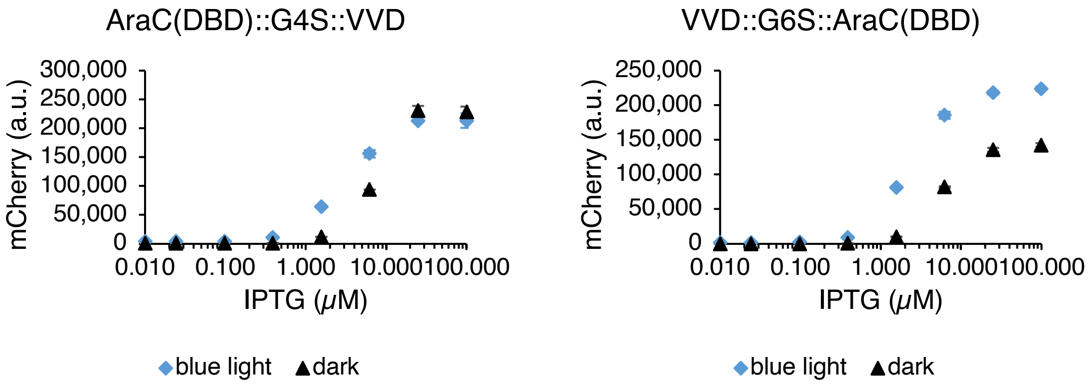


**Supplementary Fig. 16| Examples of IPTG-dose response curves for the VVD-AraC chimeric constructs in presence of the synthetic $I_1$-$I_1$ promoter.** IPTG-dose response curves of cells transformed with the single plasmid bearing the indicated VVD-AraC fusions and mCherry under control of the synthetic $I_1$-$I_1$ promoter. Cells were either in the dark, or induced with saturating blue light. Values represent mean ± s.d. of either triplicates or duplicates for mCherry fluorescence.

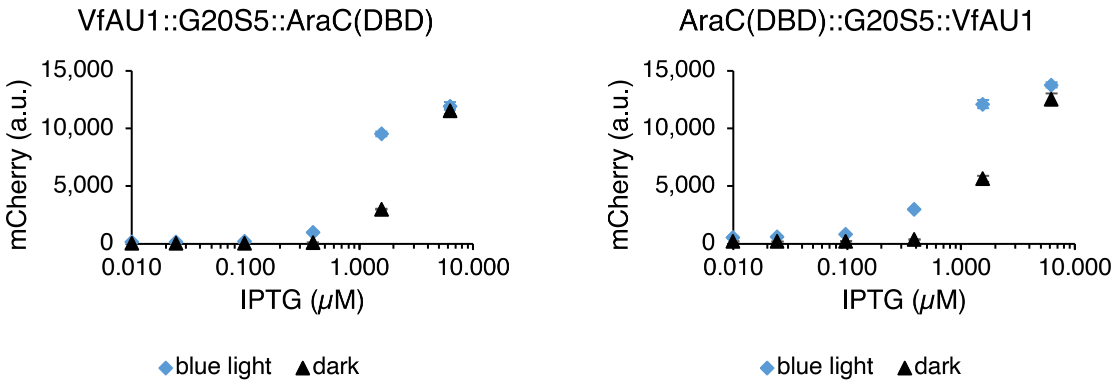


**Supplementary Fig. 17| Examples of IPTG-dose response curves for the VfAu1-AraC chimeric constructs in presence of the synthetic $I_1$-$I_1$ promoter.** IPTG-dose response curves of cells transformed with the single plasmid bearing the indicated VfAu1-AraC fusions and mCherry under control of the synthetic $I_1$-$I_1$ promoter. VfAu1 contained the RBS RiboJ+B0033m, while VfAu1::G20S5::AraC(DBD) contained the RBS 116. Cells were either in the dark, or induced with saturating blue light. Values represent mean ± s.d. of duplicates for mCherry fluorescence.

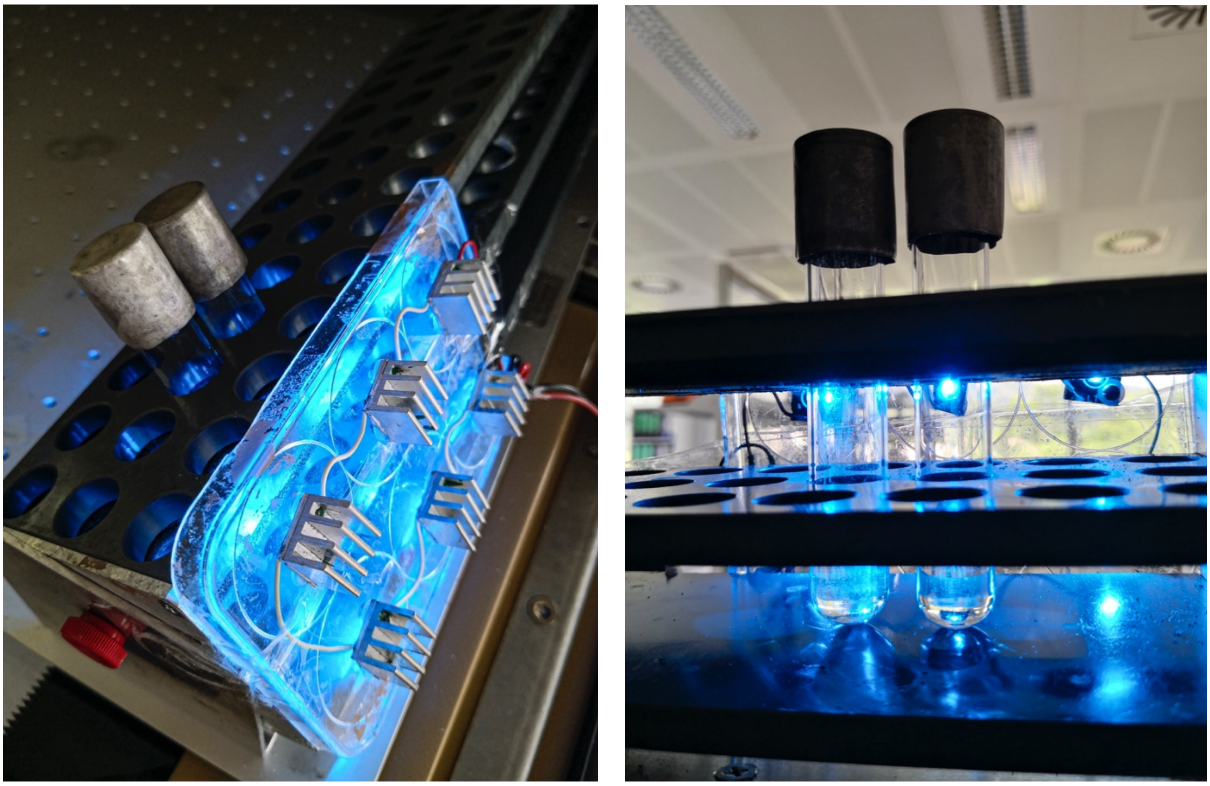

**Supplementary Fig. 18| Setup used to illuminate individual tubes in the incubator.**

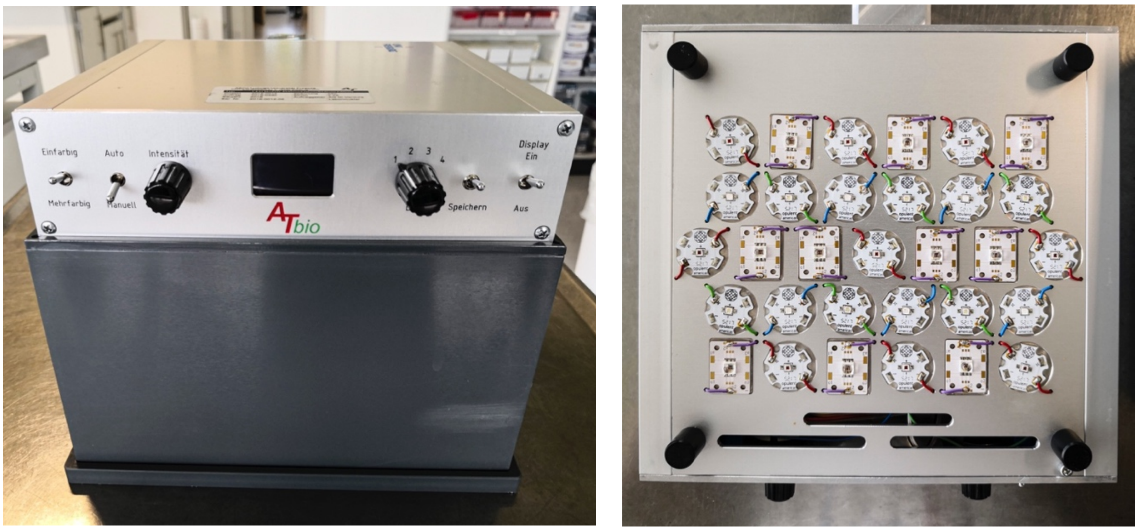


**Supplementary Fig. 19| Custom-made lightbox used to illuminate 96-well plates.** Left, front view. Right, bottom view.

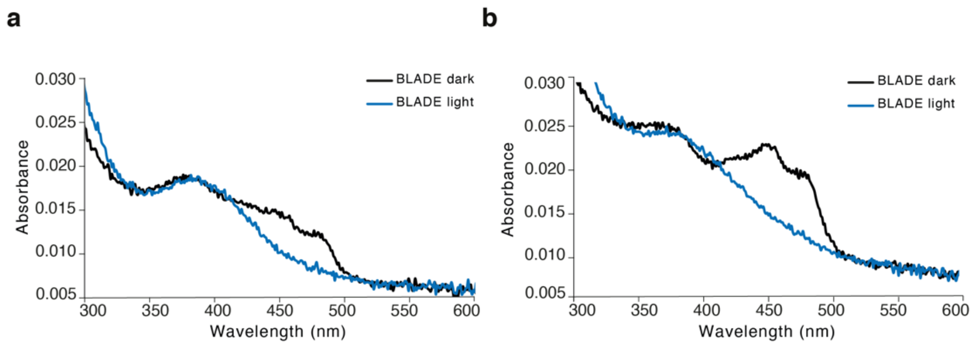


**Supplementary Fig. 20| Absorption spectrum of the FAD cofactor within BLADE FP6. a,** Absorption spectrum measured with the protein incubated 1 day in the dark. **b,** Absorption spectrum measured with the protein incubated 4 days in the dark.

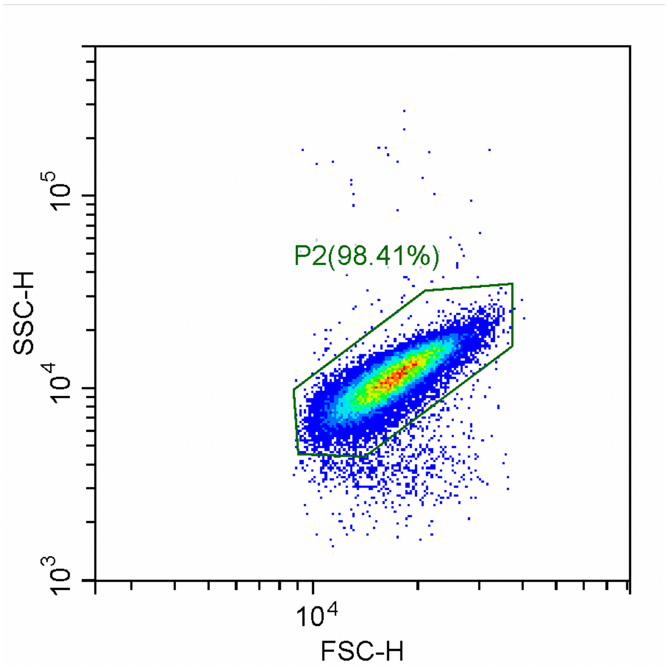


**Supplementary Fig. 21| Flow cytometry gating strategies.** Gate used for the analysis of flow cytometric data shown in Fig. 6 and Supplementary Fig. 12, 14-17. The SSC-H and FSC-H hexagon gate was drawn by eye on the AB360 control in the CytExpert v.2.1.0.92 and kept constant for all experiments using the same cell type. Shown is a screenshot of the gate taken directly from the CytExpert software.

### References


1. http://parts.igem.org/Promoters/Catalog/Anderson.
